# Tomato leaf transcriptomic changes promoted by long-term water scarcity stress can be largely prevented by a fungal-based biostimulant

**DOI:** 10.1101/2025.06.26.661737

**Authors:** Lidia López-Serrano, Alberto Férez-Gómez, Remedios Romero-Aranda, Emilio Jaime Fernández, Jesús Leal-López, Edurne Baroja-Férnandez, Goizeder Almagro, Karel Doležal, Ondrej Novak, Luis Díaz, Rocío Bautista, Rafael Jorge León Morcillo, Javier Pozueta-Romero

## Abstract

Water availability is by far the leading environmental factor limiting crop productivity worldwide. The use of cell-free microbial culture filtrates (CF) as biostimulant is gaining ground as a safe and ecologically sound approach to improving crop yields while reducing anthropogenic pressure. However, their action mechanisms remain unknown. Here we found that foliar application of *Trichoderma harzianum* CF enhanced fruit yield, root growth and photosynthesis in plants of a commercial tomato cultivar grown under long-term water deficit in Mediterranean greenhouse conditions. To investigate the biochemical and molecular mechanisms underlying this phenomenon, we adopted an integrative and systems biology approach to characterize plants grown under optimal and suboptimal irrigation conditions (OIC and SOIC, respectively) with or without the fungal CF treatment. Water shortage promoted changes in the levels of drought stress-related signalling molecules, and in the transcriptome of leaves that potentially accounted for the physiochemical differences recorded between OIC– and SOIC-grown plants. Notably, many of these changes were largely prevented by the foliar application of fungal CF to SOIC-grown plants, including the expression of ca. 50% of the expression of water scarcity-responsive genes. These genes did not respond to CF in OIC-grown plants, indicating that the transcriptomic response to CF is strongly determined by the water status of the plant. Taken together, the data provided evidence that foliar application of fungal CF improves yield and long-term water scarcity tolerance by preventing a large portion of the stress-induced transcriptional response, rendering plants “blind”, or less sensitive to water stress.

## INTRODUCTION

Water availability is by far the leading environmental factor that limits the global productivity of major crops by directly reducing plant potential yield, but also by indirectly influencing their interactions with biotic factors that play a critical role in the world’s food security, especially in semi-arid and dry regions (Lesk et al., 2016). Plants have evolved a plethora of morphological, physiological, biochemical and molecular adaptations to overcome the detrimental effects caused by water deficit, including enhanced photosynthesis, activation of antioxidative defence mechanisms, accumulation of osmoprotectants, modification of the ion, metabolic and hormonal balances, transcriptome reprograming, etc. In addition to the problems derived from limited water availability, the environmental damage inflicted by practices based on depletion of soil and water resources and intensive application of fertilizers has become a major limitation in conventional agriculture. Therefore, it is mandatory to establish new strategies for a sustainable and eco-friendly agriculture aimed at enhancing crop’s yield, water scarcity tolerance and water use efficiency, while reducing the negative impact of agrochemicals on the environment. Over the last years, compiling evidence has shown that many of these objectives can be achieved by the use of biostimulants (Benito et al., 2024; Cerruti et al., 2024).

A common and environmentally friendly cultural practice to increase crop yield and/or protect plants from different stresses is based on the soil inoculation with plant growth promoting microorganisms as biostimulants. However, this technology faces a number of challenges, including inconsistences in soil chemical and physical properties, specific crop’s growth requirements and host compatibility (Morcillo et al., 2022). Microorganisms emit diffusible compounds that promote root branching and nutrient uptake, improve photosynthesis, enhance tolerance to (a)biotic stresses, and stimulate resident beneficial microbial communities in the phytosphere, thereby boosting plant growth and yield (De-la-Peña and Loyola-Vargas, 2014; Camarena-Pozos et al., 2019; Hacquard and Martin, 2024). Consistently, the application of these compounds either in a pure form or in cell-free microbial culture filtrates (CF) enhances crop yield and stress tolerance, which could be a step to minimize shortfalls related to the technology of microbial inoculation (Morcillo et al., 2022). However, studies on the biochemical and molecular mechanisms involved in the response of plants to microbial CF have not yet been carried out. Recently, we showed that soil application of cell-free CF obtained from different beneficial and pathogenic fungi promoted root growth and enhanced yield in pepper plants grown under optimal irrigation conditions (OIC) (Baroja-Fernández et al., 2021). This response was associated with strong proliferation of the resident beneficial soil microbiota (Baroja-Fernández et al., 2021). We thus hypothesized that the positive effect exerted by this treatment on growth and yield is at least partly due to mechanisms involving activation of the beneficial soil microbiota. Fungal CF contain volatile organic compounds (VOC) that, once distilled and applied to soil, promoted responses similar to those triggered by the fungal CF (Baroja-Fernández et al., 2021). Therefore, volatile constituents of CF have strong action potential and act as major determinants of the response of plants to soil application of fungal CF.

In Southern Mediterranean countries of Europe where water scarcity is already a limiting factor for agricultural production, the expected droughts exacerbated by climate change will lead to significant reductions of water resources for intensive tomato (*Solanum lycopersicum*) greenhouse production (Silva et al., 2017). Numerous studies have been conducted to investigate the adaptations of different tomato cultivars to water deficit (Zhou et al., 2019; Liu et al., 2023; Shu et al., 2024). Furthermore, recent reports have provided valuable insights into the molecular action mechanisms of different biostimulants to mitigate the harmful consequences of this stress in tomato (Hamedeh et al., 2022; Lucia et al., 2022; Cerruti et al., 2024). However, most of these studies were carried out using non-commercial tomato cultivars grown under highly controlled environmental conditions in growth chambers and, in some of them, effects on fruit yield were not characterized. In addition, while transcriptional changes promoted by the biostimulants were reported, these were not compared with those promoted by water scarcity. Therefore, it was not possible to evaluate whether the action mechanisms of the bioestimulants were due to the stimulation of new processes that confer tolerance to water deficit or to mitigation/prevention of the transcriptional response triggered by this stress. To decipher the biochemical and molecular mechanisms underlying the mode of action of fungal CF, here we have adopted an integrative approach, from a yield evaluation, to a metabolic, hormonal and transcriptional characterization of plants of a commercial tomato cultivar grown in a semi-commercial greenhouse under OIC and suboptimal irrigation conditions (SOIC), with or without application of cell-free CF of the beneficial fungus *Trichoderma harzianum*. To minimize direct effects on plant-microbe interactions in the rhizosphere and changes in the soil microbiota, the fungal CF was applied as foliar spray. The results presented in this work highlight novel regulatory mechanisms of the responses of plants to long-term water deficit and provide a comprehensive overview of novel, transcriptionally regulated action mechanisms of a fungal-based biostimulant, offering valuable information for sustainable agriculture.

## RESULTS

### Tomato fruit yield and composition

Water scarcity exerted a negative effect on number, size and yield of commercial fruits (**Figure 1**). Foliar application of three CF dilutions (D1, D2 and D3) increased by 20% and 11% the total number and yield of commercial fruits in SOIC-grown plants in the 2021 and 2022 campaigns, respectively (**Figure 1, Supplemental Figure 1**). These treatments did not alter significantly the mean fruit weight in SOIC-grown plants (**Figure 1**). In SOIC-grown plants, water input required to obtain the same amount of fruits was ca. 20% lower than in OIC-grown plants and was further reduced by the CF treatment (**Supplemental Figure 2**). Fruits of SOIC-grown plants accumulated more soluble sugars and less organic acids than those of OIC-grown plants (**Supplemental Figure 3A,B**). The CF treatment enhanced the levels of sugars and total carotenoids in fruits of OIC-grown plants and, in general, those of organic acids in fruits of plants grown under the two irrigation conditions (**Supplemental Figure 3**). Moreover, the CF treatment promoted ionomic changes in fruits of OIC-grown plants (i.e., it enhanced the levels of Mn, Fe, Cu, Zn, Na^+^, P, and reduced those of Cl^−^, PO_4_^3-^, NO_3_^−^ and SO_4_^3-^), but had no effect in those of SOIC-grown plants **(Supplemental Table 1).**

**Figure 1:**
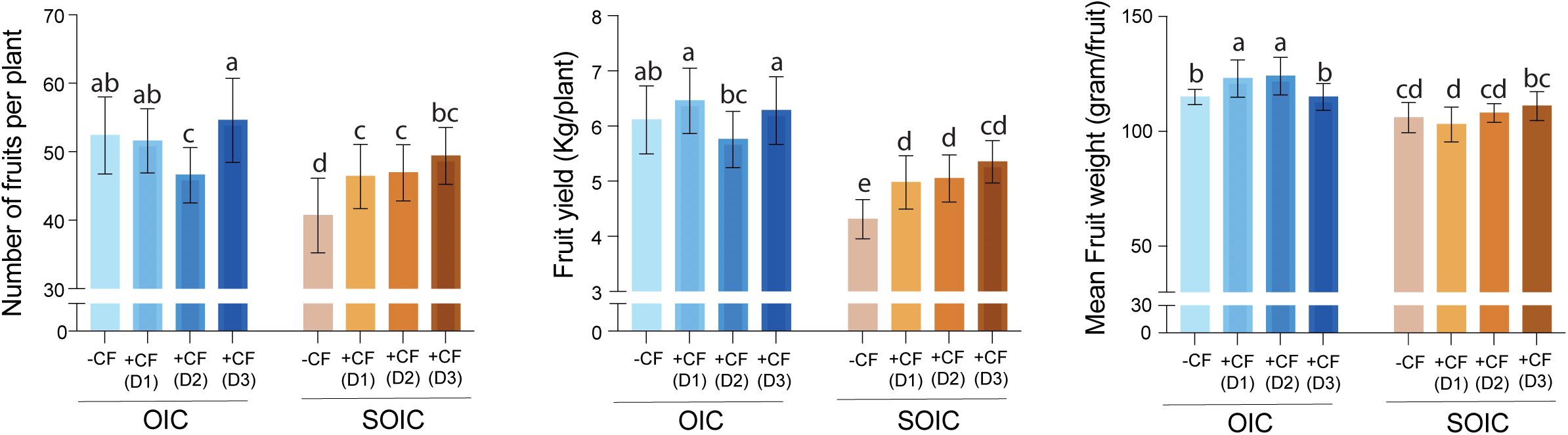
Effect of foliar application of fungal CF on tomato fruit production. Number, yield and mean weight of commercial fruits of tomato plants grown in the 2021 campaign under optimal or suboptimal irrigation conditions (OIC and SOIC, respectively) and with or without foliar application of D1, D2 and D3 fungal CF dilutions. Values represent the means ± standard deviations (SD) obtained from 10 plants. Different letters indicate significant differences at P < 0.05 (LSD test).

### Plant growth and photosynthesis

We further evaluated the effectivity of the CF treatment by measuring the leaf, stem and root dry weights (DW) in OIC– and SOIC-grown plants at the end of the experiment. Furthermore, we recorded visible symptoms of wilting during the central hours of the day, and measured *A_n_* and plant height. As shown in **Supplemental Figure 4A**, vegetative biomass production of SOIC-grown plants was lower than that of OIC-grown plants. In SOIC-grown plants, the CF treatment had no significant effects on shoot biomass, but enhanced root growth (**Supplemental Figure 4A**). In addition, this treatment enhanced *A_n_* in SOIC-grown plants (**Supplemental Figure 4B**). SOIC-grown plants were ca. 30% shorter than OIC-grown plants (**Supplemental Figure 5A**). This difference in plant height between OIC– and SOIC-grown plants was significantly reduced by the CF treatment (**Supplemental Figure 5A**). Under the highest evaporative water demand conditions, SOIC-grown plants exhibited wilting leaves, which could be mitigated by the CF treatment (**Supplemental Figure 5B**).

### Effect of foliar application of CF on leaf composition

#### Primary metabolism

Leaves of OIC– and SOIC-grown plants exhibited different amino acid content profiles (**Supplemental Table 2**). Under reduced water supply conditions, plants accumulated higher levels of the glutamate (Glu)-derived, stress-related amino acids gamma-aminobutyric acid (GABA), histidine (His), proline (Pro) and aspartate (Asp) than those of OIC-grown plants (**Figure 2A**). In leaves of plants grown under the two irrigation conditions, foliar CF application enhanced His levels and reduced those of Pro. Furthermore, this treatment enhanced GABA levels in leaves of OIC-grown plants, and those of Glu and Asp in leaves of SOIC-grown plants (**Figure 2A**). Leaves of SOIC-grown plants accumulated lower levels of starch and higher levels of soluble sugars (e.g. glucose and fructose) than those of OIC-grown plants, but differences were reduced by the CF treatment on SOIC-grown plants (**Figure 2B**).

**Figure 2:**
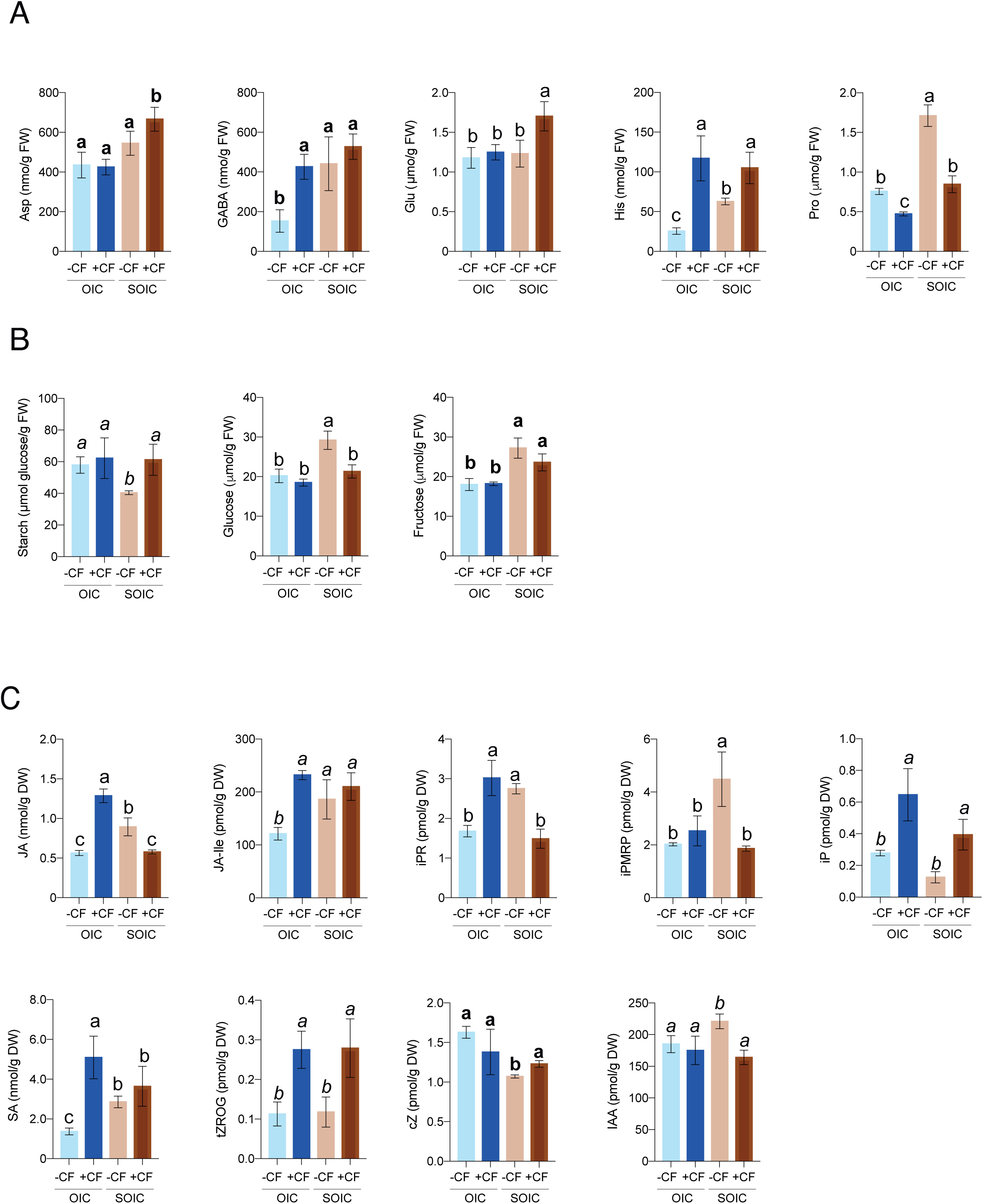
Effect of foliar application of fungal CF on tomato leaf composition. Contents of. (A) amino acids, (B) soluble sugars and (C) hormones in leaves of tomato plants grown in the 2021 campaign under optimal or suboptimal irrigation conditions (OIC and SOIC, respectively), with or without foliar CF (D3) application. Different letters indicate significant differences at P < 0.05 (LSD test). Normal letters indicate the LSD of the interaction (Treatment x Irrigation), while *italic* letters (Treatment) and **bold** letters (Irrigation) were used for the LSD of the main factors when the treatment-irrigation interaction was not significant. Values represent the means ± SD obtained from 4 replicates of 10 plants. Further information about leaf composition can be found in **Supplemental Tables 3** and **4**.

#### Phytohormone content

The hormonomes of leaves of OIC– and SOIC-grown plants were different (**Supplemental Table 3** and **Supplemental Figure 6)**. Compared with leaves of OIC-grown plants, those of plants grown under reduced water supply conditions accumulated lower levels of the main active cytokinin (CK) forms isopentenyladenine (iP) and cis-zeatin (cZ), and higher levels of iP precursor and transport forms (e.g. iPR and iPRMP), free jasmonic acid (JA) and conjugated with isoleucine (JA-Ile), salicylic acid (SA) and indole acetic acid (IAA) (**Supplemental Table 3, Figure 2C**). Foliar CF application altered the overall leaf hormonome (**Supplemental Table 3, Supplemental Figure 6**). In the two irrigation conditions, this treatment enhanced the levels of iP and those of the *trans*-zeatin (tZ) O-glucoside storage form tZROG (**Supplemental Table 3, Figure 2C**). Some of the hormonal changes promoted by the CF treatment depended on the water status of the plant. Thus, unlike in OIC-grown plants, the treatment enhanced the cZ content and reduced those of IAA, iPR, iPRMP and JA in leaves of SOIC-grown plants (**Supplemental Table 3, Figure 2C**). Furthermore, unlike in SOIC-grown plants, foliar CF application augmented the levels of JA, JA-Ile, iPR and SA (**Supplemental Table 3, Figure 2C**).

#### Mineral content

Long-term water scarcity and CF treatment promoted changes in the mineral composition of leaves. Some of these ions play important roles in respiration, photosynthesis and protection against oxidative stress (e.g. Cu^2+^, Fe^2+^) (Cai et al., 2024), amino acid metabolism (e.g. NO_3_^−^ and SO_4_^2-^), fruit carbohydrate metabolism (e.g. Cl^−^) (Wan et al., 2025) and osmotic adjustment for drought stress alleviation (e.g. Cl^−^ and Na^+^) (Erel et al., 2014; Franco-Navarro et al., 2015). As shown in **Supplemental Table 4**, leaves of SOIC-grown plants accumulated higher levels of Cl^−^

, Na^+^, Cu^2+^ and Mo, and lower levels of NO_3_ ^−^ than those of OIC-grown plants. In the two irrigation conditions, the CF treatment enhanced the levels of Cu^2+^ and reduced those of Cl^−^ (**Supplemental Table 4**). Also, the treatment enhanced the levels of NO_3_^−^ and SO_4_^2-^ in leaves of SOIC-grown plants and those of Zn and Fe^2+^ in leaves OIC-grown plants (**Supplemental Table 4**). Moreover, it reduced leaf Na^+^ content in SOIC-grown plants.

### Principal component and heat map analyses of data

Principal component (PC) and heatmap analyses using Pearson hierarchical clustering of the data for all evaluated parameters were performed to investigate the mathematical distribution of the replicates in terms of dependence on the irrigation condition and foliar CF application treatment. The results obtained confirmed that the irrigation condition and the CF treatment were important determinants of the physiological and biochemical properties of the plant. The distribution of the data in both fruits and leaves showed a clustering between the four growth conditions, both in the PC (**Figure 3A**) and heatmap (**Figure 3B**) analyses. The PC analyses revealed that the clusters of the CF treatment in OIC– and SOIC-grown plants were distanced from those of the non-treated plants (**Figure 3A**). Notably, the CF treatment brought the cluster of fruits of SOIC-grown plants closer to that of fruits of OIC-grown plants (**Figure 3A**), strongly indicating that, in terms of fruit production and quality, the CF treatment made SOIC-grown plants more “OIC-like”.

**Figure 3:**
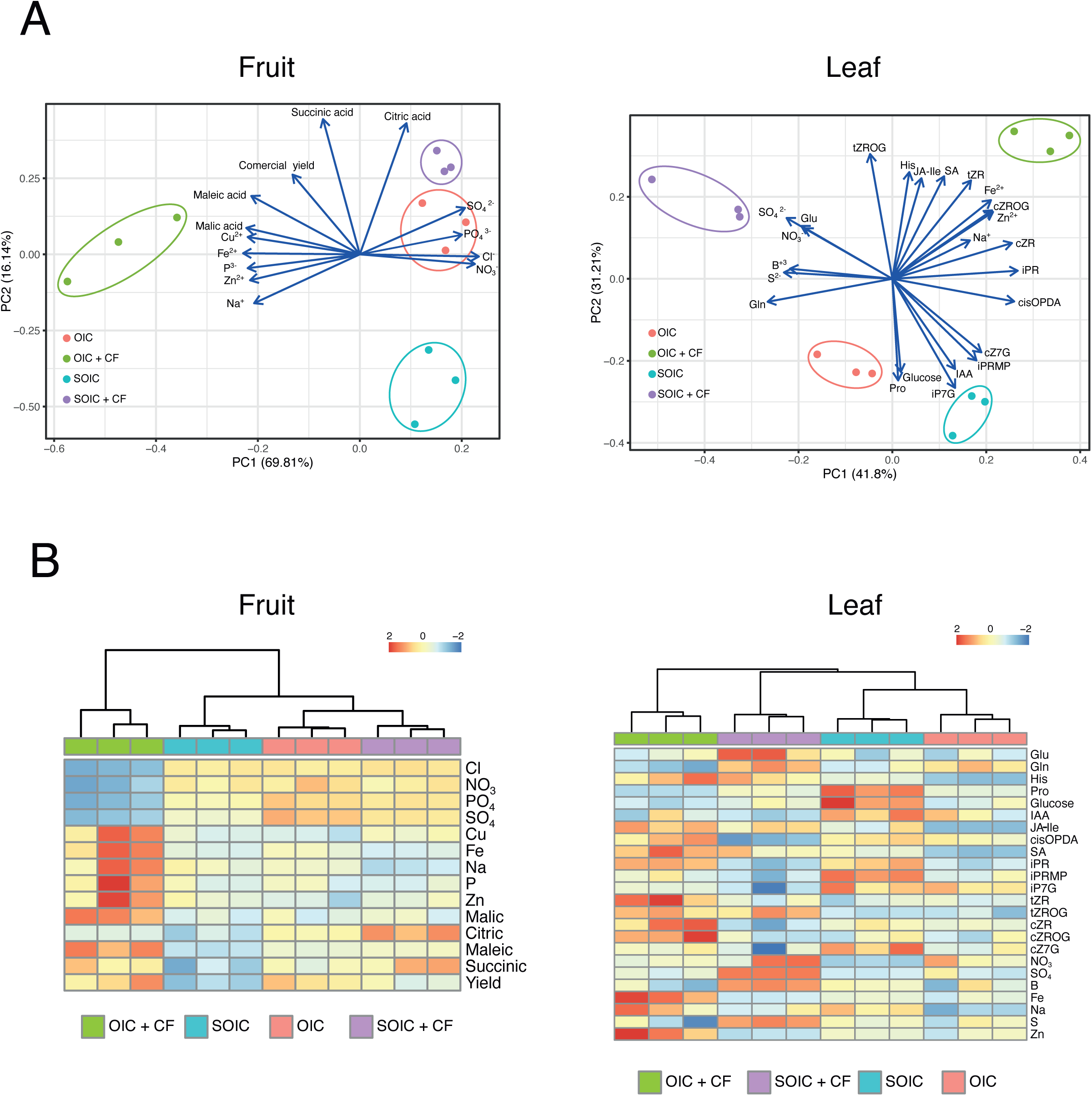
Principal component analyses and heatmap of the physiological and biochemical parameters in fruits and leaves of plants grown under optimal and suboptimal irrigation conditions. (A, B) Biplot of the principal component analysis of the distribution of replicates and (C, D) clustered heatmap of the correlation of the treatments with respect to the physiological and biochemical parameters.

### Effect of foliar application of CF on the leaf transcriptome

To investigate the molecular basis of the response to long-term water deficit stress and biostimulant treatment, we performed high-throughput transcriptome analyses of leaves of OIC– and SOIC-grown plants with or without foliar application of fungal D3 CF dilution. Quantitative real-time RT-qPCR analyses of some of the identified genes not only validated the results of this study (**Supplemental Figure 7**), but also showed that foliar application of the D2 CF dilution promoted transcriptomic changes similar to those promoted by D3 application (**Supplemental Figure 8**). As shown in **Supplemental Table 5** and **Figure 4**, these analyses revealed that, compared with leaves of OIC-grown plants, 575 out of the 20759 transcripts identified in this study were up-regulated and 689 genes were down-regulated in leaves of SOIC-grown plants. Studies on the biological processes affected by the irrigation regime using the MapMan tool revealed that these 1264 “drought responsive genes” are involved in multiple processes including photosynthesis, primary (e.g. carbohydrate, amino acid, lipid, nucleotide, etc.) metabolism, secondary metabolism, phytohormone synthesis and signalling, redox homeostasis, RNA biosynthesis and processing, protein synthesis, homeostasis and modification, solute transport, etc. (**Supplemental Table 5, Figure 4**). In leaves of SOIC-grown plants, the CF treatment enhanced the levels of 1698 transcripts and reduced those of 1566 (**Supplemental Table 6, Figure 5A**). Furthermore, it enhanced the levels of 1008 transcripts and reduced those of 479 in leaves of OIC-grown plants (**Supplemental Table 7, Figure 5B**). Among the 1487 genes differentially expressed by the CF treatment in leaves of OIC-grown plants, 288 (hereinafter designated as “core CF responsive genes”) were also differentially expressed by the CF treatment in leaves of SOIC-grown plants (**Supplemental Table 8, Figure 5C,D**). Notably, 243 out of the 575 transcripts that were upregulated by water scarcity, and 395 out of the 689 transcripts that were downregulated by this stress, exhibited opposite regulation upon CF treatment in leaves of SOIC-grown plants (**Supplemental Table 9, Figure 6A,B**). Consistently, t-SNE analyses of the transcriptomic data revealed that the CF treatment brought the cluster of SOIC-grown plants closer to that of OIC-grown plants (**Figure 6C**). We refer to these 638 genes whose response to long-term water scarcity can be largely prevented or mitigated by the CF treatment as the “golden genes”.

**Figure 4:**
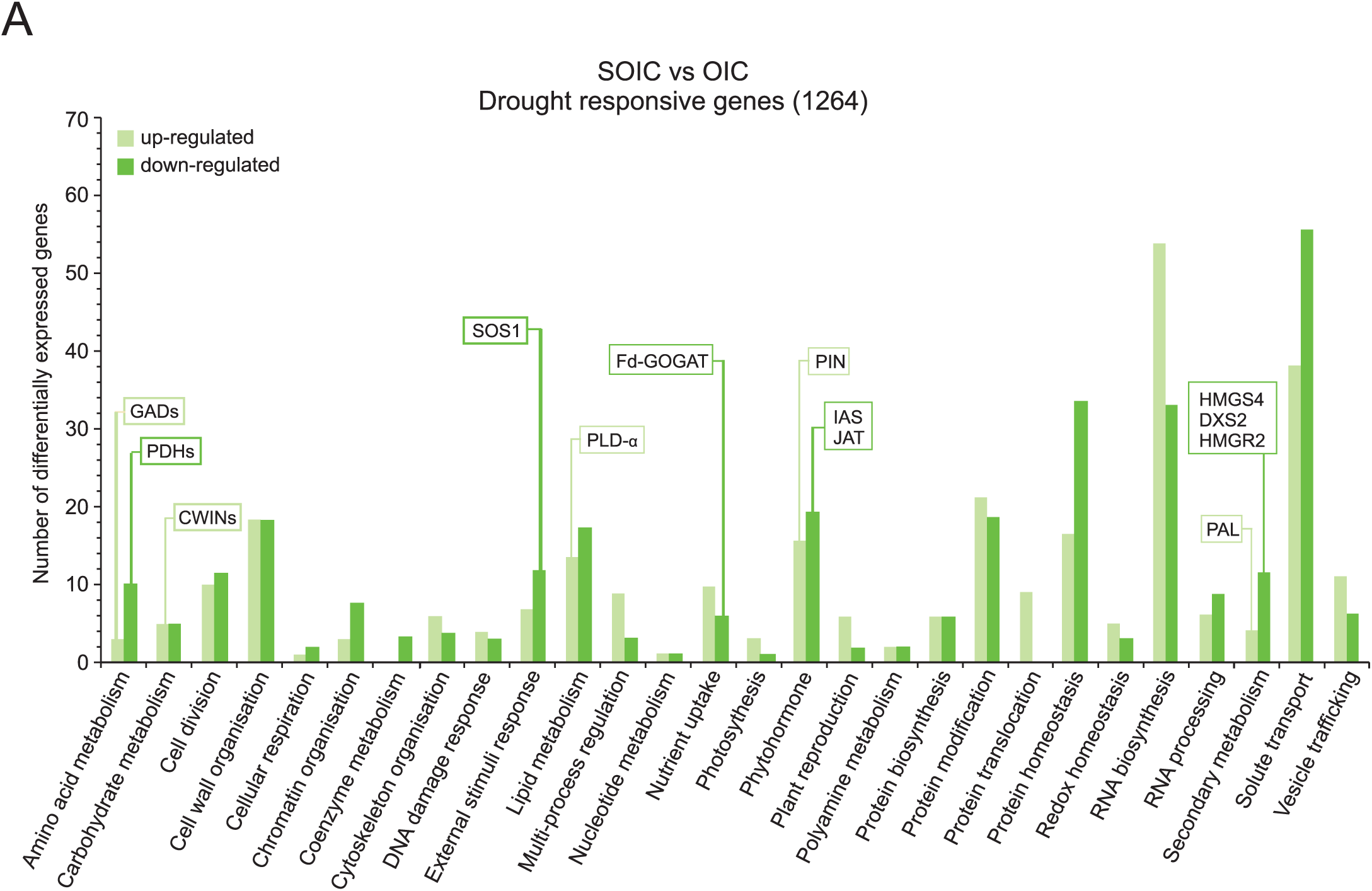
Transcriptome profiling of leaves of tomato plants grown under optimal and suboptimal irrigation conditions. Functional categorization of the transcripts differentially expressed in leaves of tomato plants grown under optimal and suboptimal irrigation conditions (OIC and SOIC, respectively) (see also **Supplemental Table 5**). Drought responsive genes were sorted according to putative functional categories assigned by MapMan software. Numbers of up– and down-regulated transcripts in each categorical group are indicated by pale and deep green bars, respectively. DEGs discussed here are boxed.

**Figure 5:**
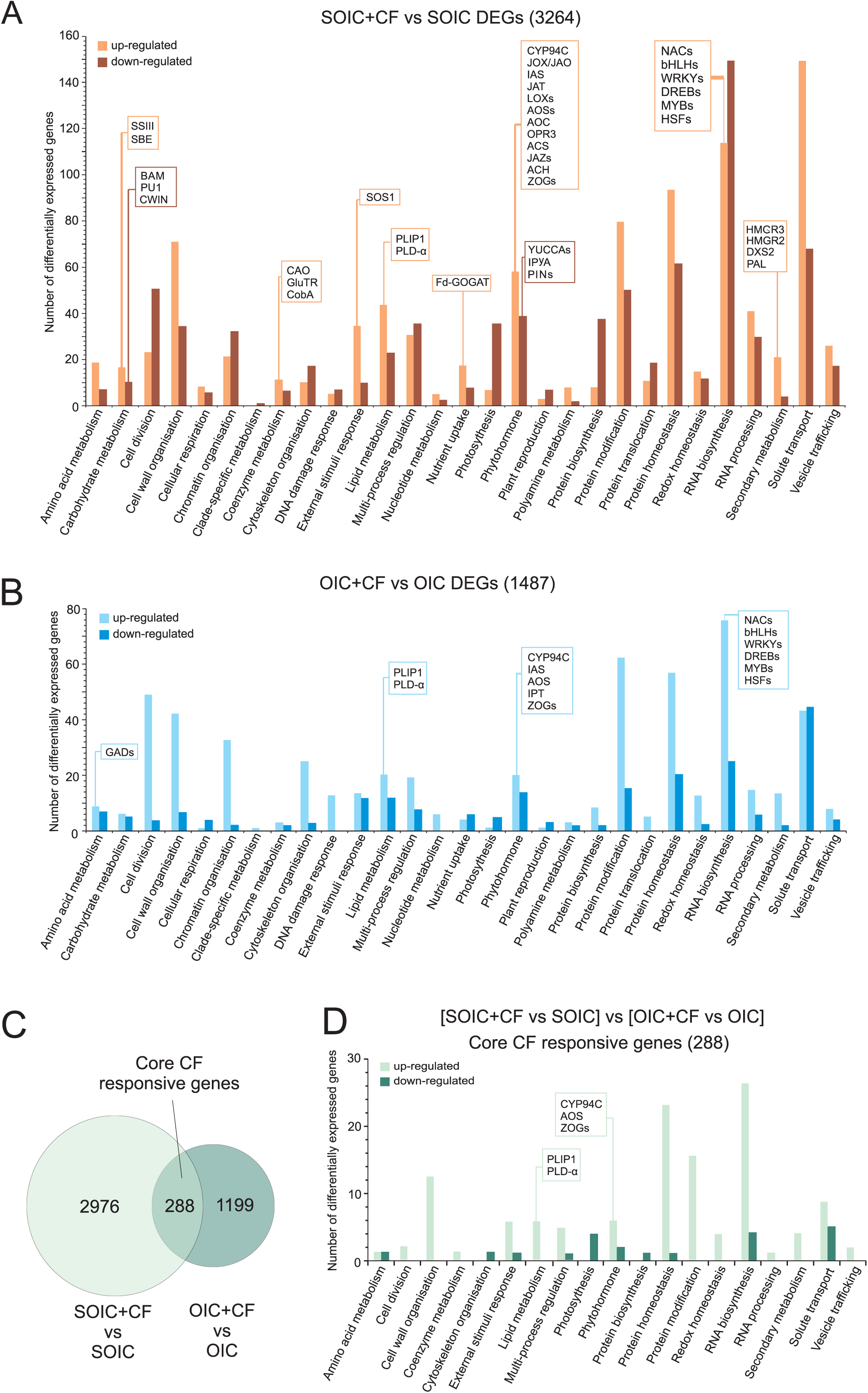
Identification of “core CF responsive genes”. (A) Functional categorization of the transcripts differentially expressed in leaves of tomato plants grown under SOIC + CF vs. SOIC conditions (see also **Supplemental Table 6**). Numbers of up– and down-regulated transcripts in each categorical group are indicated by pale and deep brown bars, respectively. (B) Functional categorization of the transcripts differentially expressed in leaves of tomato plants grown under OIC + CF vs. OIC conditions (see also **Supplemental Table 7**). Numbers of up– and down-regulated transcripts in each categorical group are indicated by pale and deep blue bars, respectively. (C) Venn diagram analysis showing DEGs of [SOIC+CF vs SOIC] and [OIC+CF vs OIC]. The 288 overlapping DEGs are the “core CF responsive genes”. (D) Functional categorization by the Mapman software of the “core CF responsive genes” (see also **Supplemental Table 8**). Numbers of up– and down-regulated transcripts in each categorical group are indicated by pale and deep green bars, respectively. DEGs discussed here are boxed.

**Figure 6:**
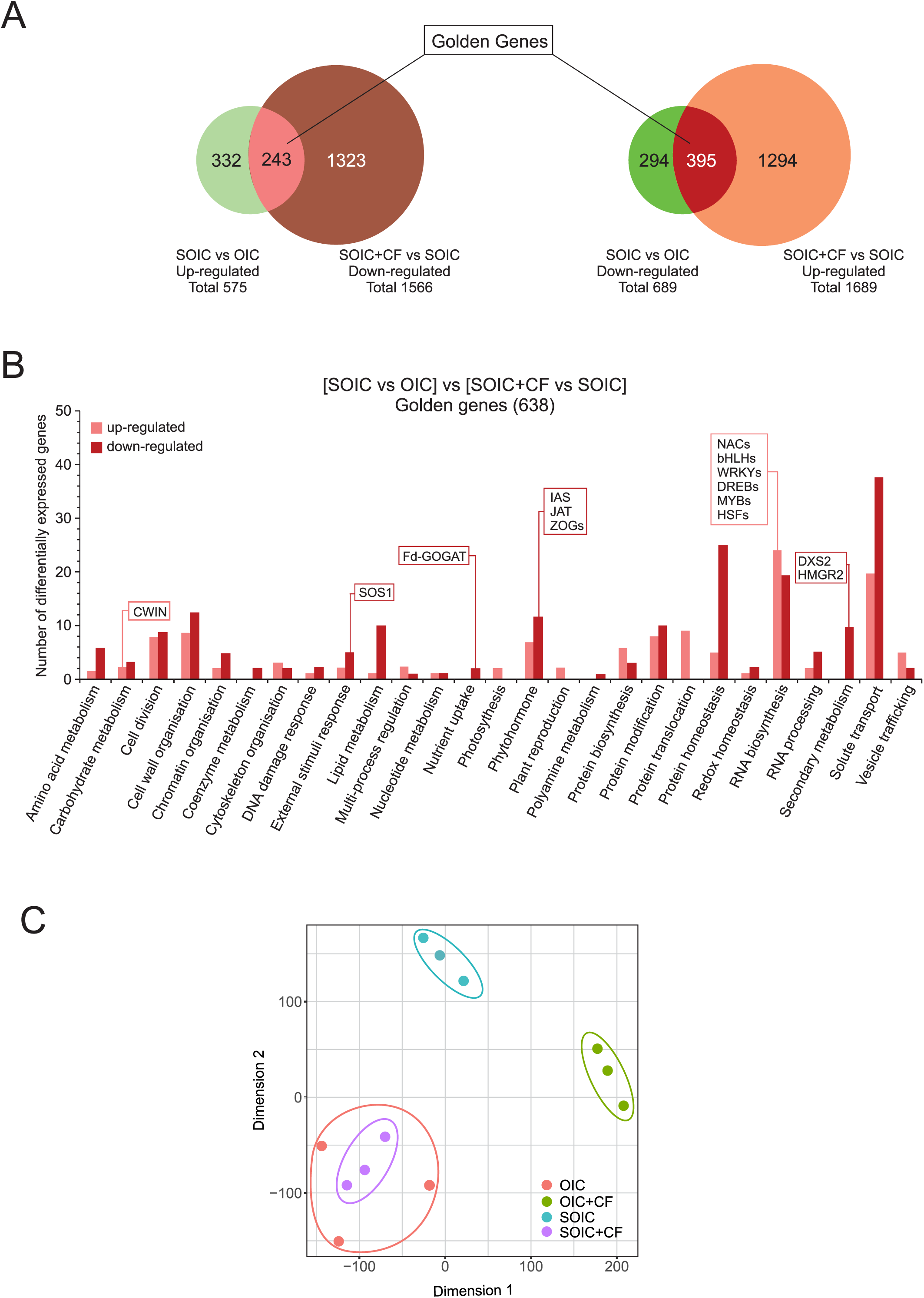
Identification of “golden genes”. (A) Venn diagram analysis illustrating the 638 golden DEGs identified by comparing water scarcity-promoted transcriptomic changes (SOIC vs OIC) with those promoted by the CF in SOIC-grown plants (SOIC+CF vs SOIC). (B) Functional categorization of the “golden genes” (see also **Supplemental Table 9**). Numbers of up– and down-regulated transcripts in each categorical group are indicated by pale and deep red bars, respectively. DEGs discussed here are boxed. (C) t-SNE plot of the RNAseq data.

## DISCUSSION

### Foliar application of fungal CF can be used as an eco-friendly practice to enhance tomato fruit yield

In a previous study we showed that soil application of cell-free fungal CF enhanced fruit yield and root growth in pepper plants grown under OIC conditions, and promoted the proliferation of beneficial soil microbiota (Baroja-Fernández et al., 2021). Therefore, we hypothesized that the effect exerted by fungal CF on fruit yield and root growth could be due, at least in part, to the activation of beneficial soil microbiota (Baroja-Fernández et al., 2021). Here we showed that foliar application of fungal CF, which minimizes direct effects on plant-microbe interactions in the rhizosphere, increased commercial fruit number and yield in SOIC-grown tomato plants (**Figure 1**). Furthermore, the CF treatment promoted fruit compositional changes, mainly in OIC-grown plants (**Figure 3, Supplemental Figure 3, Supplemental Table 1)**. This would indicate that the mechanisms triggered by soil and foliar applications of fungal CF are different, at least partly. We also showed that foliar CF application reduced the water footprint (**Supplemental Figure 2**), enhanced root growth and photosynthesis (**Supplemental Figure 4**), and reduced temporary symptoms of long-term water deficit stress (**Supplemental Figure 5**) in SOIC-grown plants. Therefore, foliar fungal CF application can be used as an eco-friendly practice to enhance tomato fruit yield under water stress conditions.

### Foliar application of CF largely prevents changes in the levels of stress-related compounds promoted by long-term water deficit

Many mechanisms developed by plants to mitigate deleterious effects infringed by water deficit are regulated by phytohormones, sugars and amino acids. In general, this stress enhances auxin, ABA, SA and JA action and reduces that of CK (Rivero et al., 2009; Kim et al., 2013; Miura and Tada, 2014; Ali et al., 2020; Casanova-Sáez et al., 2022). Water scarcity also reduces leaf starch levels and enhances those of soluble sugars (mainly glucose) and Glu-derived amino acids (e.g. Pro, GABA and His) (Zanella et al., 2016; Thalmann and Santelia, 2017). These compounds play important functions in osmotic protection, ROS scavenging, control of hormone homeostasis and signaling, gene expression regulation, etc. (Moore et al., 2003; Ruan et al., 2010; Sairanen et al., 2012; Kushwah and Laxmi, 2014; Mekonnen et al., 2016; Qiu et al., 2020; Ji et al., 2022). In addition, water scarcity promotes the accumulation of Cl^−^ and Na^+^ to reduce osmotic potential (Erel et al., 2014; Franco-Navarro et al., 2015), and sustain water and nutrient use efficiencies (Franco-Navarro et al., 2019) and photosynthetic capacity. Consistently, leaves of SOIC-grown plants accumulated lower levels of starch, iP and cZ, and higher levels of IAA, JA, JA-Ile, SA, GABA, His, Asp, Pro, soluble sugars, Cl^−^ and Na^+^ than those of OIC-grown plants (**Figure 2, Supplemental Tables 2, 3** and **4**). Furthermore, leaves of SOIC-grown plants accumulated higher levels of iPR and iPRMP than those of OIC-grown plants, which would indicate that long-term water deficit stress exerts a negative effect on the conversion of iPR and iPRMP to iP.

Results presented here showing that foliar CF treatment enhanced fruit yield and root growth, and altered fruit composition in SOIC-grown plants would strongly indicate that the fungal CF have constituents (mainly VOC, Baroja-Fernández et al., 2021) that may trigger changes in the balance of stress– and signalling-related compounds in leaves. Consistent with this idea, the CF treatment reduced IAA and Pro levels, and enhanced those of His, iP and tZROG in leaves of both SOIC– and OIC-grown plants (**Figure 2, Supplemental Tables 2** and **3**). Furthermore, unlike in SOIC-grown plants, the CF treatment enhanced the levels of GABA, JA, JA-Ile, iPR and SA in leaves of OIC-grown plants. Moreover, the treatment promoted the accumulation of Asp and Glu and the reduction of the levels of soluble sugars (mainly glucose), iPR, iPRMP and JA in leaves of SOIC-grown plants (**Figure 2**). Notably, levels of Pro, starch, glucose, IAA, iP, iPR, iPRMP and JA in leaves of SOIC-grown plants treated with CF were comparable to those of OIC-grown plants (**Figure 2**). The overall data would strongly indicate that in SOIC-grown plants, foliar application of CF largely prevented changes in the levels of stress-related signals thereby reducing stress perception and rendering them more “OIC-grown like” plants and more tolerant to long-term water deficit. This action mechanism is different to those of other biostimulants which promote the accumulation of stress-related signals (e.g. glucose, IAA and JA) in tomato plants (Ertani et al., 2017; Casadesús et al., 2019). The differences in the responses of stress– and signalling-related compounds to the CF treatment in SOIC– and OIC-grown plants would indicate that the effect of the CF treatment is strongly determined by the water status of the plant.

### Physiochemical changes promoted by long-term water deficit stress can be explained by transcriptome reprogramming

Many transcriptomic differences in leaves of OIC– and SOIC-grown plants involved transcripts coding for proteins that catalyze rate-limiting steps of transcriptionally regulated metabolic pathways involved in drought tolerance. Thus, leaves of SOIC-grown plants accumulated higher levels of transcripts that code for phospholipase D-α (PLD-α) and phenylalanine ammonia lyase (PAL), and lower levels of IAA-amido synthetase (IAS) transcripts than those of OIC-grown plants (**Supplemental Table 5, Figure 4**). PLD-α is involved in the early steps of JA biosynthesis (Wang et al., 2018), whereas PAL catalyzes the first committed regulatory step of one main pathway of SA biosynthesis (Miura and Tada, 2014). IAS catalyzes the rate-limiting step of synthesis of IAA-amino acid conjugates, providing a mechanism for the plant to reduce pools of active IAA, maintain IAA homeostasis and enhance drought tolerance (Ding et al., 2008; Casanova-Sáez et al., 2022). Therefore, the high IAA, JA, Ile-JA and SA levels in leaves of SOIC-grown plants (**Figure 2C**) may be due, at least in part, to reduced IAS and enhanced PLD-α and PAL activities, respectively.

Leaves of SOIC-grown plants accumulated lower levels of transcripts coding for enzymes that catalyze main rate regulatory steps of the plastidial methylerythritol-4-phosphate (MEP) pathway (e.g. the isoform 2 of deoxy-D-xylulose 5-phosphate synthase (DXS2)) and the cytosolic mevalonate (MVA) pathway (e.g. the isoform 4 of 3-hydroxy-3-methylglutaryl-CoA synthase (HMGS4) and the isoform 2 of 3-hydroxy-3-methylglutaryl-CoA reductase (HMGR2)) than those of OIC-grown plants (**Supplemental Table 5, Figure 4**). Both pathways feed the synthesis of isoprenoid compounds that play crucial roles in plant growth, development, regulation of gene expression, metabolism, etc. (Estévez et al., 2001; Suzuki et al., 2004; Córdoba et al., 2009; Paetzold et al., 2010). In addition, the MEP pathway functions as a stress sensor that negatively regulates IAA biosynthesis and transport (Jiang et al., 2018). Therefore, the high IAA levels in leaves and the reduced growth and biomass production of SOIC-grown plants can be due, at least partly, to reduced activities of the MEP and MVA pathways.

GABA and Pro levels are largely determined by the drought inducible Glu decarboxylase (GAD) and the catabolic enzyme Pro dehydrogenase (PDH), respectively (Kiyosue et al., 1996; Shelp et al., 1999). Pro levels are also negatively regulated by SOS1, a plasma membrane Na^+^/H^+^ antiporter that plays a prime role in Na^+^ homeostasis and osmotic adjustment in response to osmotic stress (Shi et al., 2003). Leaves of SOIC-grown plants accumulated higher levels of transcripts coding for GAD and lower levels of PDH and SOS1 encoding transcripts than those of OIC-grown plants (**Supplemental Table 5, Figure 4**). Therefore, high levels of Pro and GABA in leaves of SOIC-grown plants can be due, at least partly, to reduced SOS1 and PDH and enhanced GAD activities.

The pool size of soluble sugars under water deficit stress is strongly determined by starch and sucrose breakdown activities (Ruan et al., 2010). Cell wall invertases (CWIN) catalyze the hydrolytic breakdown of sucrose into glucose and fructose. Soluble sugars negatively regulate CK signaling (Kushwah and Laxmi, 2014), and positively regulate IAA signaling (Sairanen et al., 2012). CWIN play important regulatory roles in CK-dependent fruit development and composition and CK-mediated leaf senescence (Lara et al., 2004; Jin et al., 2009). Leaves of SOIC-grown plants accumulated higher levels of CWIN encoding transcripts than those of OIC-grown plants (**Supplemental Table 5, Figure 4**), which could account for the high levels of glucose and IAA, and low levels of active CK in leaves of SOIC-grown plants.

We compared our RNAseq dataset with previous gene expression analyses related to water deficit stress in tomato plants. Among the 220 and 3765 drought-responsive genes identified in the Jinlingmeiyu and M82 cultivars grown in chambers under highly controlled conditions (Zhou et al., 2019; Li et al., 2023), 3 and 197 were present in the list of 1264 “drought responsive genes” identified in the present study, respectively. Moreover, among the 332 core drought-responsive genes identified in the LA1375– and Moneymaker-1 cultivars grown in greenhouses (Shu et al., 2024), 37 were present in our list of “drought responsive genes”. It thus appears that the molecular mechanisms involved in the water scarcity stress response is highly genotype-dependent and strongly influenced by developmental stage, stress intensity and environmental growth conditions.

### Physiochemical changes promoted by the foliar CF application can be explained by transcriptome reprogramming

Foliar CF application induced significant transcriptomic changes in leaves of SOIC– and OIC-grown plants (**Supplemental Tables 6** and **7**, **Figure 5**), which further provides evidence that CF constituents have strong action potential. Many of these changes differed between SOIC– and OIC-grown plants, further indicating that the effect of the CF treatment is strongly determined by the water status of the plant. This treatment enhanced the expression of several transcription factors (TF) of the NAC, bHLH, WRKY, DREB, MYB and HSF families (**Figure 5A,B**), some of which directly involved in the responses to different stress types and drought tolerance (Sakuma et al., 2006; Wu et al., 2012; Bai et al., 2018; Liu et al. 2024). In leaves of SOIC-grown plants, the CF treatment upregulated the levels of transcripts coding for starch synthesis enzymes (e.g. SSIII and SBE), and downregulated those coding for starch and sucrose breakdown enzymes (e.g. BAM, PU1 and CWIN) (**Supplemental Table 6, Figure 5A**). Moreover, this treatment enhanced the levels of transcripts coding for MEP pathway enzymes (e.g. DXS2, HMGR2 and HMGR3) (**Supplemental Table 6, Figure 5A**). These changes can potentially account for the CF-promoted accumulation of starch and reduction of soluble sugars in leaves (**Figure 2**), and enhancement of growth and biomass in SOIC-grown plants (**Supplemental Figure 5A, Supplemental Figure 4A**). In leaves of OIC– and SOIC-grown plants, the CF treatment enhanced the expression of genes coding for PAL and various *trans*-zeatin O-glucosyltransferases (ZOG) involved in the synthesis of O-glucoside active CK reserves and that of CK synthesis enzymes (e.g. IPT) (**Supplemental Tables 6 and 7**, **Figure 5A,B**). In addition, the treatment downregulated the expression of IAA biosynthesis genes (e.g. IPyA and YUCCA) and upregulated those of the IAA-inactivating IAS and numerous proteins involved in JA biosynthesis (e.g. PLD-α, phosphatidylglycerol lipase [PLIP1], CYP94C, JAO, AOS, LOX, ACS, AOC, OPR3 and ACH) (**Supplemental Tables 6 and 7**, **Figure 5A,B**). These transcriptomic changes can potentially explain the CF-promoted enhancement of SA, iP, *t*ZROG and Ile-JA levels and reduction of IAA levels in leaves of both OIC– and SOIC-grown plants (**Figure 2C, Supplemental Table 3**). In leaves of SOIC-grown plants, the CF treatment upregulated the levels of transcripts coding for SOS1 and the Glu biosynthesis limiting enzyme Fd-GOGAT (**Supplemental Table 6, Figure 5A**). Therefore, it is likely that the reduced Pro levels and enhanced levels of Glu and the Glu-derived His and Asp promoted by the CF treatment are due to upregulation of Fd-GOGAT and SOS1 expression. In addition, in leaves of OIC-grown plants, the CF upregulated GAD expression (**Supplemental Table 7, Figure 5B**), which could account for the high GABA levels.

To further explore the molecular basis of the responses to the CF treatment, we compared our RNAseq dataset with that of a recent study related to the effect of foliar application of ERANTHIS® in OIC– and SOIC-grown Micro Tom tomato plants (Cerruti et al., 2024). ERANTHIS® is a biostimulant derived from the algae *Ascophyllum nodosum* and *Laminaria digitata* and yeast extracts. We found that 65 out of the 135 transcripts up-regulated by ERANTHIS® in SOIC-grown plants were also up-regulated by the fungal CF (**Supplemental Table 10**). Only 2 of these 65 transcripts up-regulated by the two biostimulants are members of the set of “golden genes” identified in this work (**Supplemental Table 10**). Furthermore, 53 out of the 172 transcripts up-regulated by ERANTHIS® in OIC-grown plants were also up-regulated by the fungal CF (**Supplemental Table 10**). It thus appears that, in SOIC-grown plants, different biostimulants can activate the expression of genes not responding to water deficit.

### Foliar application of CF operates by preventing or mitigating changes in the expression of a large pool of drought responsive genes in SOIC grown plants

The transcriptional response of tomato plants to water deficit identified in this work involved changes in the expression of two sets of “drought responsive genes”: (i) a set of 626 genes whose expression is not affected by the CF treatment in SOIC-grown plants, which included genes that code for various isoforms of PDH, CWIN, GAD, PAL and PLD-α (**Supplemental Tables 5** and **6**) and whose response could at least partly explain the high levels of Pro, glucose, GABA, His, SA and Ile-JA in leaves of SOIC-grown plants (**Figure 2**); and (ii) a set of 638 “golden genes” whose transcriptional response to water deficit is largely mitigated or prevented by the CF treatment in leaves of SOIC-grown plants (**Supplemental Table 9, Figure 6**). This list includes genes that code for proteins such as DXS2, HMGR2, CWIN, IAS, Fd-GOGAT and SOS1 whose expression could at least partly account for the low levels of iP and cZ, and high levels of glucose, IAA, Pro, Na^+^ and stress-related amino acids (e.g. Asp, His and GABA) in leaves of SOIC-grown plants (**Figure 2, Supplemental Table 4**). This strongly indicated that foliar CF application to SOIC-grown plants largely operated by preventing or mitigating changes in the expression of many “drought responsive genes”, rendering plants “blind” or less sensitive to stress, and more transcriptionally similar to “OIC-grown like” plants. To our knowledge, this is the first report providing evidence for the prevention or mitigation of the expression of a massive number of “drought responsive genes” as a possible mechanism of a microbial-based biostimulant to enhance drought tolerance and yield.

We also identified a set of 288 “core responsive CF genes” that are differentially expressed by CF in leaves of both OIC– and SOIC-grown plants (**Supplemental Table 8, Figure 6D**). Most of these genes were upregulated by the CF treatment, especially genes encoding TF and genes related with lipid metabolism, phytohormone action, protein modification and homeostasis and cell wall organization (**Supplemental Table 8, Figure 5D**). This list of “core responsive genes” includes PLD-α, PLIP1, CYP94C, AOS and ZOGs encoding genes whose expression could at least partly account for the enhanced tZROG and JA-Ile contents promoted by the CF treatment in leaves of both OIC– and SOIC-grown plants. When comparing the lists of “core responsive CF genes” and “golden genes”, we found that less than 4% of the 638 “golden genes” are “core responsive CF genes”. This strongly indicated that (i) CF-promoted changes in the expression of “golden genes” only occur under drought stress conditions and (ii) the transcriptomic response to CF is strongly determined by the physiological state of the plant.

### Concluding remarks

In this study, we provided valuable information indicating that foliar CF application to SOIC-grown plants works by preventing global changes in the levels of water deficit-responsive signalling molecules and “golden” transcripts, thereby reducing stress perception and increasing plant tolerance to water deficit stress. Further efforts are required to investigate whether the molecular mechanisms involved in mitigating the stress response promoted by CF also extend to other layers of regulation including the translational and post-translational ones. We propose that the identification of “golden genes” by comparing the transcriptional responses triggered by different stress types with those triggered by the application of biostimulants to plants subjected to the same stress is a valid approach to (i) elucidate the molecular mechanisms by which biostimulants confer stress tolerance, (ii) identify biomarkers to monitor stress tolerance conferred by biostimulants, and (iii) propose eco-sustainable and environmentally friendly strategies to increase stress tolerance in crops based on the manipulation of the stress-responsive gene expression.

## MATERIALS AND METHODS

### Experimental site and greenhouse conditions

Assays were conducted in two spring-summer campaigns (2021 and 2022) in a polyethylene greenhouse located at the “Instituto de Hortofruticultura Subtropical y Mediterránea” in Algarrobo (Málaga, Spain; 36°45’32.0”N 4°02’28.7”W). No greenhouse air conditioning systems were used. During the 2021 campaign, maximum midday greenhouse air temperature ranged from 26 to 30°C, with a minimum night temperature of 18°C (**Supplemental Figure 9**). Minimum relative humidity, recorded at midday, ranged from 34 to 60%. (**Supplemental Figure 9**). Values of photosynthetic active radiation (PAR, 400-700 nm) recorded at midday were 752 ± 155 µmol m^−2^ s^−1^. During the 2022 campaign, the environmental conditions inside the greenhouse were: a maximum PAR of 494 ± 190 µmol m^−2^ s^−1^, a maximum and minimum temperature of 30 ± 4 and 15 ± 4 °C, respectively, and a maximum and minimum relative humidity of 82 ± 7 and 44 ± 10 %, respectively (**Supplemental Figure 9**).

### Plant material and growth conditions

This study was conducted using the commercial tomato variety Macizo because of its performance, fruit quality, and high demand for truss tomatoes in national and international markets. Seeds were sown in vermiculite and seedlings at the third true leaf stage were transplanted in 17 L pots filled with a mixture of peat: coconut fibre: vermiculite (45:45:10), and grown under OIC. The pots were placed on troughs with 33 cm between pots and 100 cm between troughs, resulting in a plant density of 3 plants per m^2^. The irrigation control head of the greenhouse was equipped with pumps, electrovalves and an automatic irrigation programmer. Plants were supplied with a nutrient solution containing NO_3_^−^ (1.27 mM), NH_3_ (3.52 mM), CaO (2.31 mM), MgO (0.74 mM), P_2_O_5_ (0.48 mM), K_2_O (2.16 mM), SO_3_ (1.11 mM), B (23.12 µM), Cu (2.36 µM), Fe (67.15 µM), Mn (36.40 µM), Mo (1.042 µM) and Zn (3.82 µM). The electrical conductivity (EC) was 1.5 dS m^−1^ and pH 6.73. Nutrient solution was supplied by one emitter per plant and at a flow rated of 2 L/h. Fourteen days after transplanting, at the stage of 6-8 true leaves, tomato plants were divided in two groups: plants grown under OIC, and plants grown under SOIC (60% of OIC irrigation) (**Supplemental Figure 10**). The frequency of irrigation and the volume of the nutrient solution supplied was scheduled considering the volumetric water content in the substrate, recorded by Teros 12 sensors (METER Group, Inc. USA) (**Supplemental Figure 11**). The volume of irrigation water was adjusted to ensure a leaching fraction sufficient to avoid the increase of salt in the substrate. EC and pH of nutrient solution and pot lixiviates were recorded weakly. Plants were trained vertically with a single stem. Pest control practices were those commonly used by growers in the area. Ten and 28 plants per irrigation condition and foliar treatment were randomly set throughout the greenhouse in the 2021 and 2022 campaigns, respectively.

### Preparation and foliar application of CF

*T. harzianum* (CECT 2413) CF were obtained according to Baroja-Fernández et al. (2021). During the 2021 campaign, the aerial parts of plants were sprayed with non-diluted, 1:2 or 1:4 dilution of CF (D1, D2 and D3, respectively). Along the 2022 campaign, plants were sprayed with D2. Controls plants were sprayed with diluted (1:6) Murashige and Skoog (MS) medium supplemented with 1.5 mM of glucose and fructose, which is the approximate concentration of both sugars in the CF (Baroja-Fernández et al., 2021). CF and diluted MS were applied in four moments (**Supplemental Figure 10**).

### Fruit production

Fully mature fruits from bunches in the stem’s positions 1 to 6 counted from the base of the plants were harvested and classified into commercial and non-commercial groups of fruits. Fruits weighing more than 50 g without symptoms of blossom end rot or any other type of abnormality were defined as “commercial”. The agronomic water use efficiency (WUEyld) was calculated as the ratio between the amount of irrigation water supplied and the total commercial fruit production per plant. Ten and 28 plants per irrigation condition and CF treatment were evaluated in 2021 and 2022 campaigns, respectively.

### Plant biomass and morphological parameters

In the 2021 campaign, shoot height was measured from the stem base up to the shoot apex. During the midday hours, when the highest radiation and temperature values were recorded inside the greenhouse, positions of leaves within the plant with clear temporary symptoms of dehydration were recorded using plants at the 29 days after water shortage (DAWS) moment. At the end of the 2022 campaign, leaves, stems and roots from 4 plants per irrigation condition and CF treatment were harvested. Roots were rinsed in tap water and blotted dry on paper towels. All the plant samples were then oven-dried at 75 °C for at least 48 h after which their dry weights were determined (**Supplemental Figure 10**).

### Determination of gas exchange parameters

In the 2021 campaign, the CO_2_ assimilation rates (*A*_n_) were determined using 4 plants per irrigation condition and CF (D1) treatment at the 63 DAWS moment. Measurements were conducted on newly expanded young leaves (the 8-9^th^ leaf from the apex) between the 12.00-13.30 pm (GMT) using a portable LI-COR 6400 infrared gas analyser (Li-Cor Inc., United States). The measurements were recorded under saturating light conditions and values of environmental parameters inside the leaf chamber were 900 µmol m^−2^ s^−1^ PAR, 31 ± 1°C air temperature, 410 ppm air CO_2_ and 40 ± 1% of relative humidity.

### Biochemical characterization

In the 2021 campaign, leaves (4^th^ leaf from the apex) and 20 fruits from 10 plants grown under OIC or SOIC with or without CF (D3) treatment were collected from plants at 49 and 89 DAWS, respectively, ground in liquid nitrogen and stored at –80 °C. Organic acids were quantified as described in Bahaji et al. (2015). Levels of soluble sugars, hormones, starch and amino acids were determined as described in Baroja-Fernández et al. (2021). Ions were quantified at the ionomic service of Centro de Edafología y Biología Aplicada del Segura (CEBAS, Murcia, Spain) by ICP emission spectrometry (iCAP 6000, Thermo Scientific. Cambridge, England). Values were obtained from 3 to 4 replicates of 10 plants per irrigation condition and CF treatment.

### Total RNA extraction and sequencing

In the 2021 campaign, total RNA was extracted from leaves of plants at the 49 DAWS moment using the TRIzol-chloroform method according to the manufacturer’s procedure (Invitrogen). RNA extracted from 3 replicates of 10 plants per irrigation condition and CF treatment were treated with RNAase free DNAase (Thermo Fisher Scientific, Cat. N° AM2238). Samples with no degradation signals in agarose gel and A260/A280 and A260/A230 > 2 coefficients in NanoDrop were sent to Sistemas Genómicos S.L. (Valencia, Spain) for paired-end RNA poliA sequencing by Illumina technology with 150 bp pared-ends reads in triplicate obtaining ca. 20 million reads for replicate. The generated reads were mapped against the latest version of the “*Solanum_lycopersicum*” genome provided by the Ensembl database (https://plants.ensembl.org/Solanum_lycopersicum/Info/Index). Low quality reads were then removed from the mapping using Picard Tools (https://broadinstitute.github.io/picard). Once the high-quality reads were selected, gene assembly, identification and quantification were carried out using htseq.

### RNA sequencing analysis

A bioinformatics workflow developed by the SCBI Bioinformatics Unit (University of Málaga) was employed to calculate gene expression levels, perform differential expression analysis, and conduct functional enrichment analyses. All processes were executed on a SUSE Linux Enterprise Server 11SP2 equipped with a Slurm workload manager and an Infiniband network (400 Gbps). The computational cluster, Picasso Cluster at SCBI, consists of 344 nodes, with a total of 40,000 cores and 180 TB of RAM, providing the necessary computational power for high-throughput analyses. The workflow began with a quality assessment of raw sequencing reads using FastQC (v0.11.9) and MultiQC (v1.11). These tools ensured that the sequencing data met the quality standards required for downstream analyses. Subsequently, raw reads were pre-processed to remove low-quality bases, adapters, contaminant sequences, and other unwanted content using Fastp and Ribodetector. Default parameters optimized for Illumina paired-end reads were applied. The cleaned reads were re-evaluated with FastQC and MultiQC to confirm the effectiveness of the pre-processing steps and ensure the absence of sequencing artefacts. Cleaned reads were aligned to the *S. lycopersicum* reference transcriptome (version SL4.0) obtained from the Sol Genomics Network (https://solgenomics.net/organism/Solanum_lycopersicum/genome) using Bowtie2 (v2.4.4). The alignment was performed with default parameters, along with the –-no-mixed and –-no-discordant options to retain only properly paired reads. The resulting SAM files were sorted and indexed using SAMtools, and transcript-level read counts were quantified using sam2counts (https://github.com/vsbuffalo/sam2counts). Differential expression analysis was carried out using DESeq2. DESeq2 normalizes the raw read counts by calculating size factors to account for differences in library sizes, estimates gene-specific dispersion parameters, and applies shrinkage to improve reliability. A negative binomial model was fitted to the normalized data, and hypothesis testing was performed using the Likelihood Ratio Test. Genes were considered differentially expressed if they met the criteria of a p-value ≤ 0.05 and an absolute fold change ≥ 1.5. Functional enrichment analyses were performed using over-representation analysis (ORA) and gene set enrichment analysis (GSEA) based on Gene Ontology (GO) terms. The R package ClusterProfiler was employed for these analyses, with statistical significance defined as an adjusted p-value < 0.05. Differentially expressed genes (DEGs) were mapped to MapMan bins for data visualization and pathway analysis (version 3.6.0). To this end, the tomato MapMan ontologies (http://www.gomapman.org/export/current/mapman/sly_SL2.40_ITAG2.3_2015-01-09_mapping.txt.tgz) were retrieved from the GO MapMan web resource and imported into the MapMan tool. All Venn diagrams wwere produced using the BioVenn web tool. The expression data of the samples were clustered using t-distributed Stochastic Neighbor Embedding (t-SNE), a nonlinear dimensionality reduction method commonly used for visualizing high-dimensional data (van der Maaten and Hinton, 2008). This approach ensures that replicates within the same condition cluster together, facilitating the assessment of sample similarity.

### Validation of RNA-Seq data by RT-qPCR analysis

Fourteen genes were selected for the validation of RNA-seq results by real time quantitative PCR (RT-qPCR). RNA (1.5 µg), was used for cDNA synthesis using polyT primers and a RevertAid First Strand cDNA Synthesis Kit (Thermo Scientific, USA) according to the manufacturer’s instructions. RT-qPCR was carried out using SsoFast EvaGreen Supermix (Bio-RAD) on a CFX96 real-time PCR detection system (Bio-RAD). Comparative threshold values were normalized to ACTIN internal control. Relative gene expression was derived by using 2-ι1CT, where ι1CT represents CT of the target gene minus CT of the reference gene *ACTIN*. RT-qPCR amplification was performed using primers listed in **Supplemental Table 11**.

### Statistical Analysis

Statistical analyses were performed using R Core Team statistical software (https://www.R-project.org) within the RStudio integrated development environment (http://www.posit.co). A two-way Analysis of Variance (ANOVA) was conducted using the aov) function from the stats package in R, where foliar CF application treatments (*Treatment, T*) and irrigation conditions (**Irrigation, I**) were the factors of the analysis. In the cases where interaction was found, a one-way ANOVA was performed by joining T and I. When double interaction was not found, a one-way ANOVA was performed taking into consideration only the significant factor. Means were compared by the Fisher’s least significant difference (LSD) test at *P<0.05* with the LSD.test function from agricolae package (https://CRAN.R-project.org/package=agricolae). Heatmaps were made using the Heat mapper web resource.

## ACKNOWLEDGEMENTS

We thank José Manuel Ramos Martín, Sara Sánchez Segovia, Gonzalo González Gil, Juan Francisco Ruiz Solanilla and Laura Frías for technical support.

## AUTHOR CONTRIBUTIONS

L L-S, A F-G, R J L M and J P-R designed the experiments and analyzed the data; L L-S, A F-G, R R-A, J L-L, E B-F, G A, K D, O N, L D, R B and R J L M performed most of the experiments and discussed the data; L L-S, A F-G and J P-R wrote the article with contributions from all the authors; J P-R conceived the project and obtained funding. All authors have read and agreed to the published version of the manuscript.

## SUPPLEMENTAL FIGURES

**Supplemental Figure 1:**
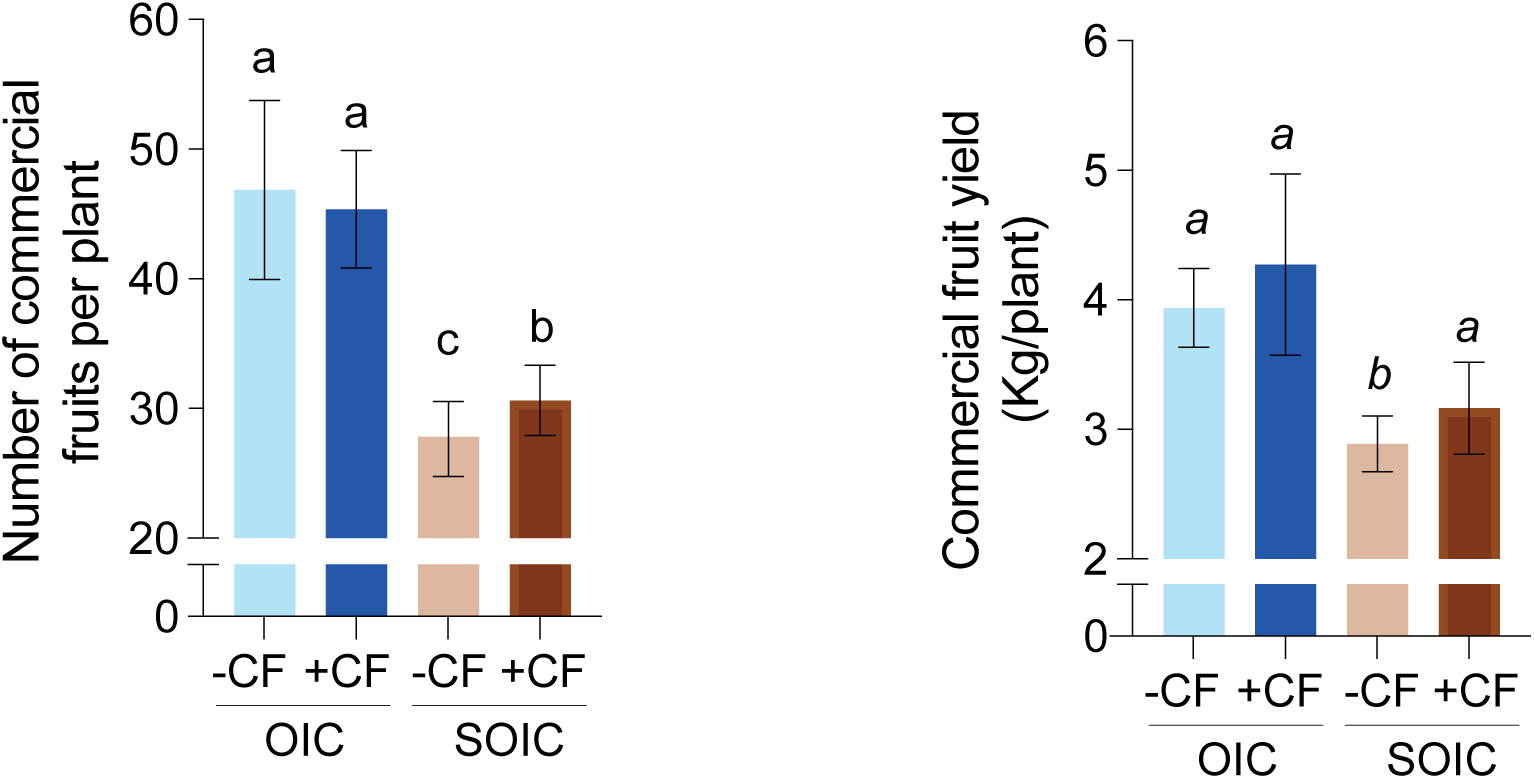
Effect of foliar application of CF on tomato fruit production. Number and yield of commercial fruits of tomato plants grown in the 2022 campaign under optimal or suboptimal irrigation conditions (OIC and SOIC, respectively) and with or without 4 foliar applications of CF (D2). Different letters indicate significant differences at P < 0.05 (LSD test). Normal letters indicate the LSD of the interaction (Treatment x Irrigation). Values represent the means ± SD obtained from 28 plants.

**Supplemental Figure 2:**
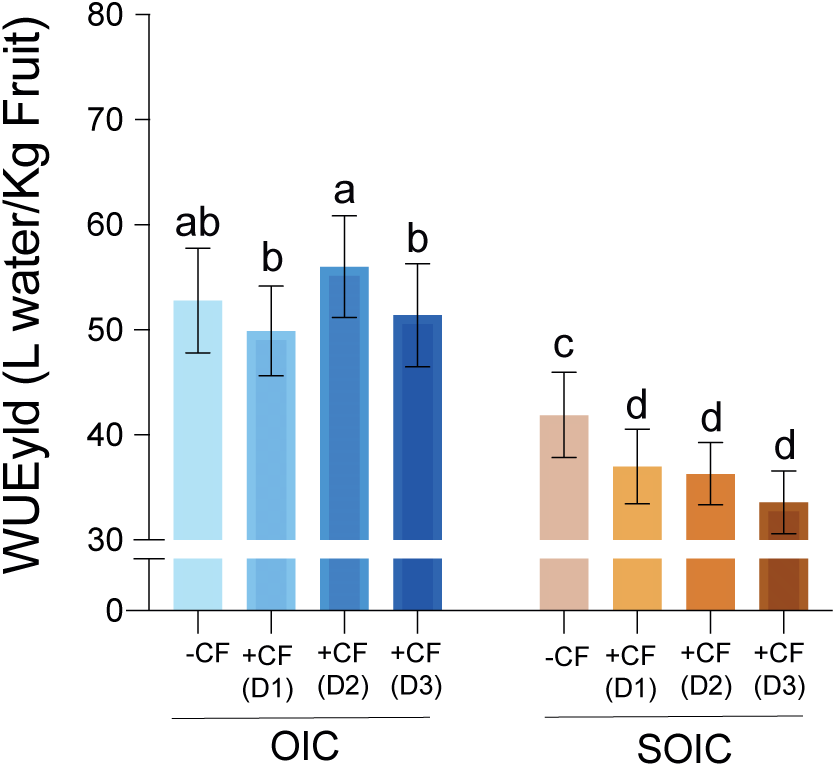
Agronomic water use efficiency (WUEyld) of tomato plants grown in the 2021 campaign under optimal or suboptimal irrigation conditions (OIC and SOIC, respectively) and with or without foliar application of D1, D2 and D3 fungal CF dilutions. Different letters indicate significant differences at P < 0.05 (LSD test). Normal letters indicate the LSD of the interaction (Treatment x Irrigation). Values represent the means ± SD obtained from 10 plants.

**Supplemental Figure 3:**
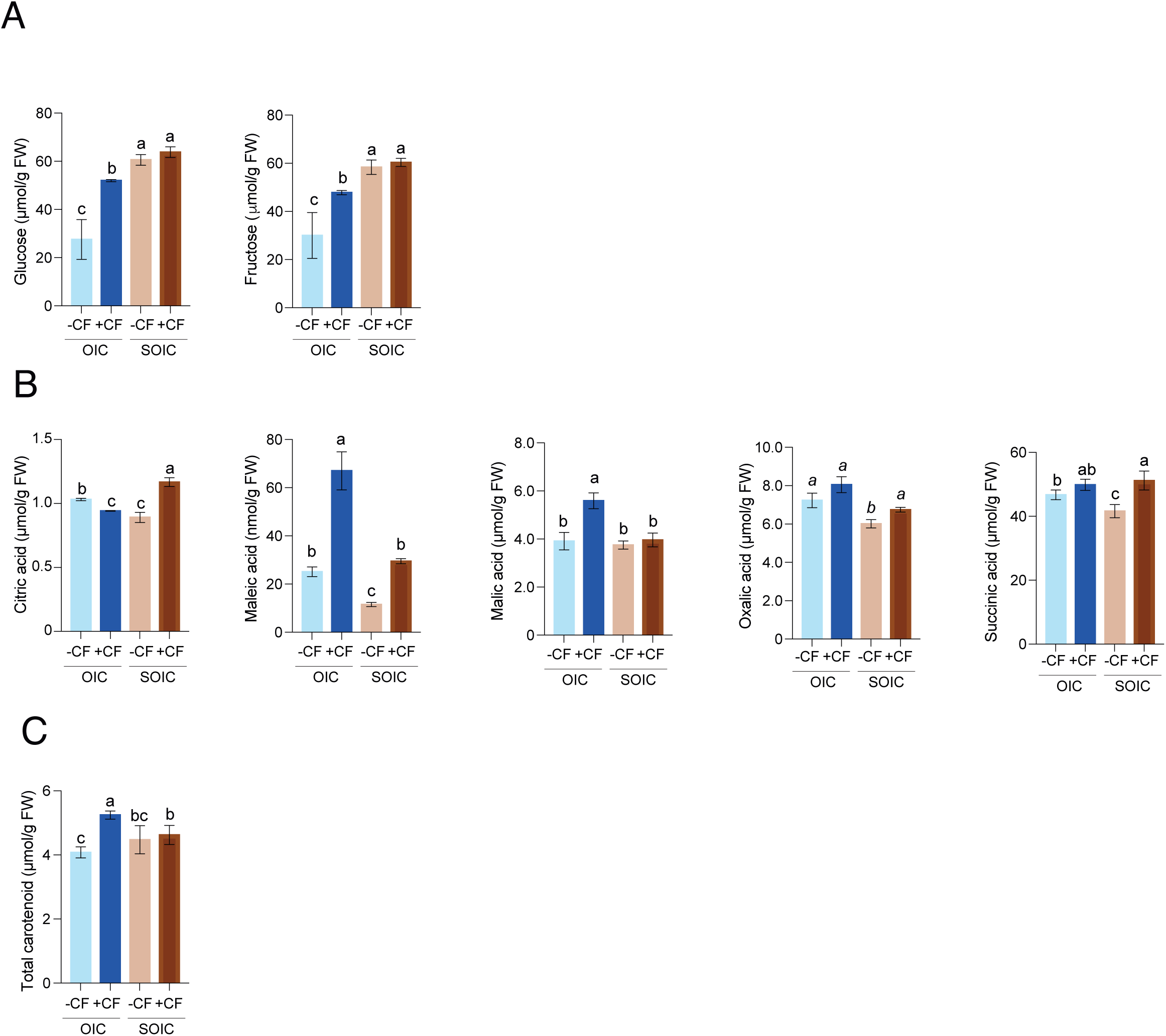
Effect of foliar application of fungal CF on tomato fruit composition. Contents of (A) soluble sugars, (B) organic acids and (C) carotenoids in fruits of tomato plants grown in the 2021 campaign under optimal or suboptimal irrigation conditions (OIC and SOIC, respectively), with or without foliar fungal CF (D3) application. Different letters indicate significant differences at P < 0.05 (LSD test). Normal letters indicate the LSD of the interaction (Treatment x Irrigation), while *italic* letters (Treatment) were used for the LSD of the main factors when the treatment-irrigation interaction was not significant. Values represent the means ± SD obtained from 4 replicates of 10 plants.

**Supplemental Figure 4:**
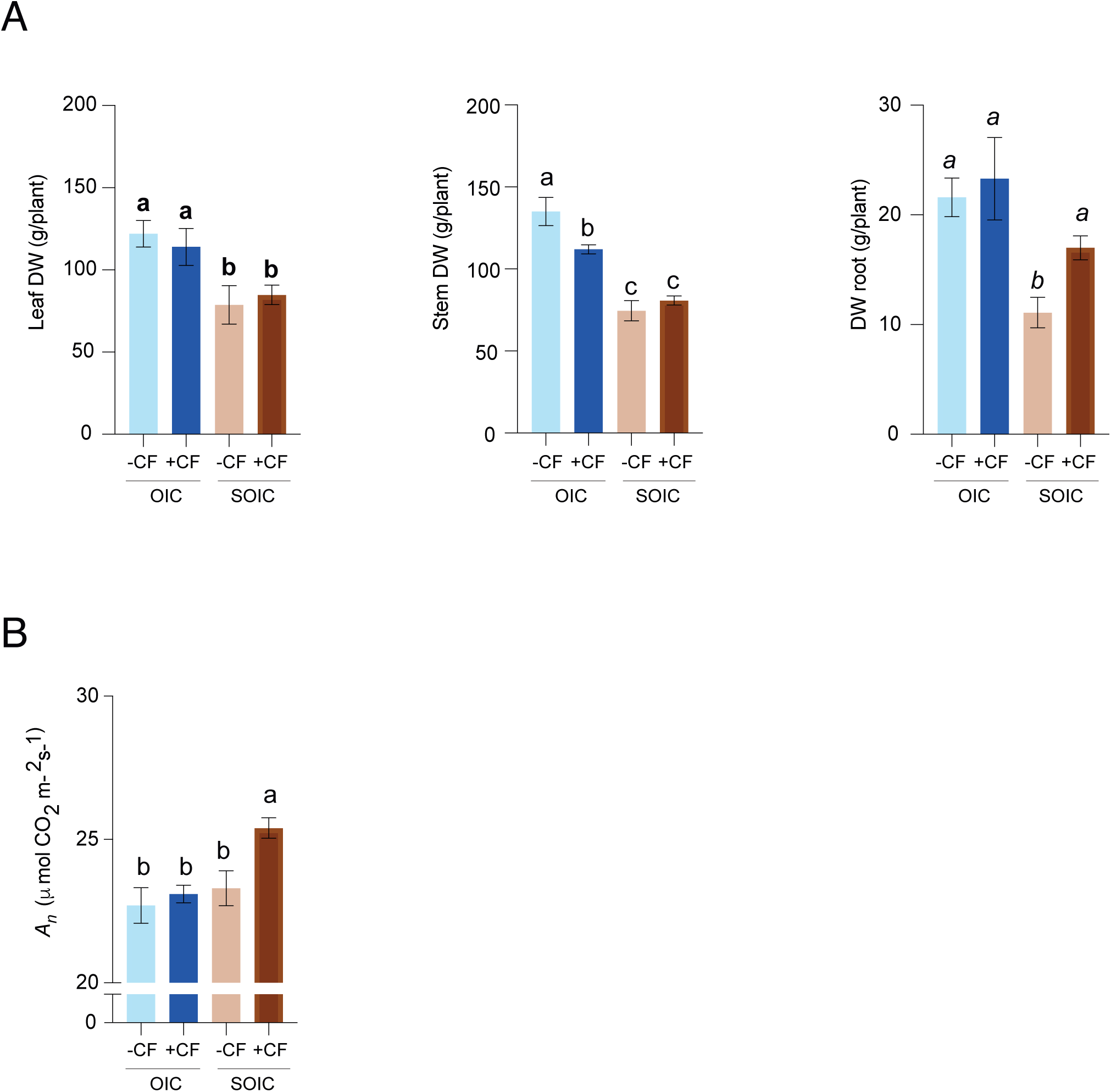
Effect of foliar application of fungal CF on plant growth and photosynthesis. (A) Leaf, stem and root dry weights (DW) and (B) CO_2_ assimilation rate (*A*_n_) of plants grown in the 2021 campaign under optimal and suboptimal irrigation conditions (OIC and SOIC, respectively), and with or without CF application. Different letters indicate significant differences at P < 0.05 (LSD test). Normal letters indicate the LSD of the interaction (Treatment x Irrigation), while *italic* letters (Treatment) and **bold** letters (Irrigation) were used for the LSD of the main factors when the treatment-irrigation interaction was not significant. Values represent the means ± standard deviations (SD) obtained from 4 plants.

**Supplemental Figure 5:**
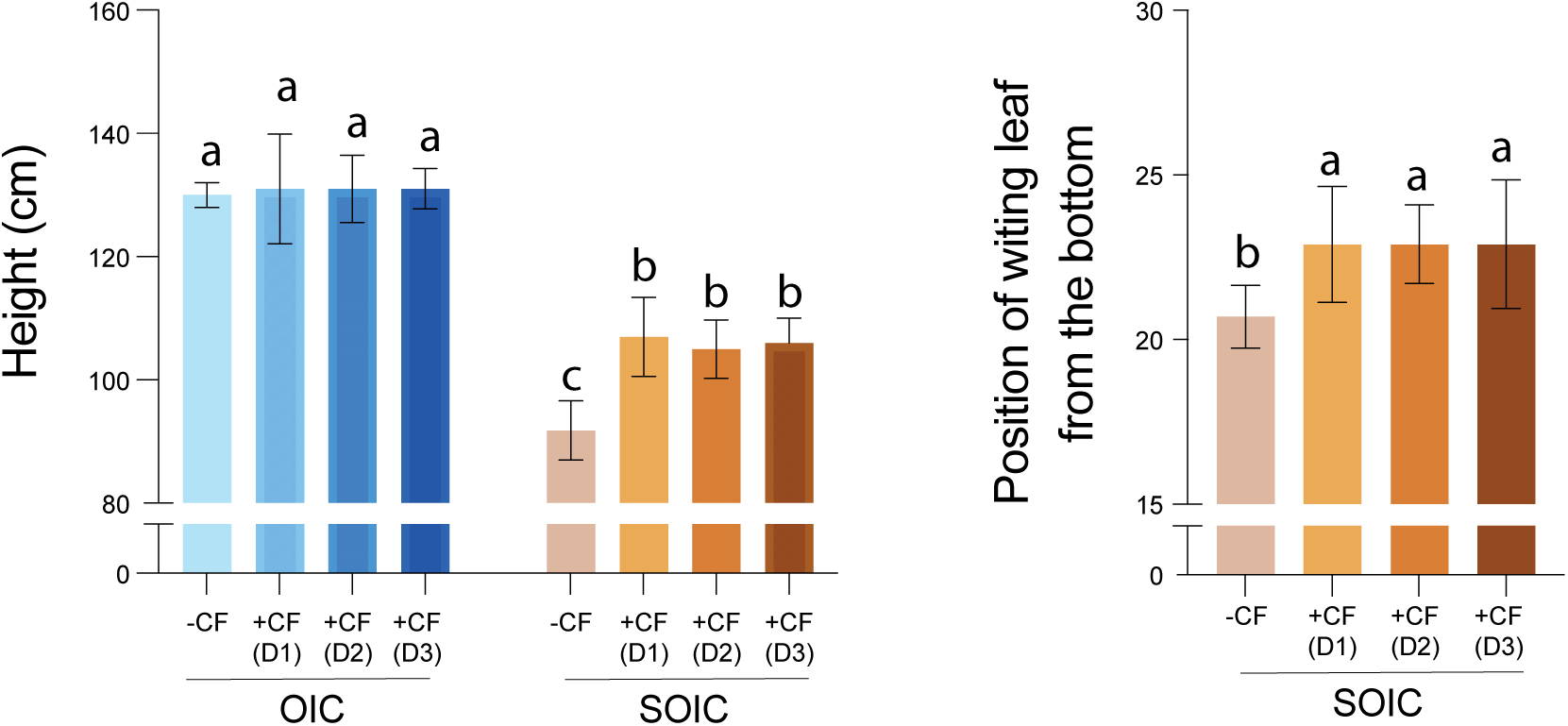
Effect of foliar application of cell-free fungal CF on morphological aspects of plants. (A) Height and (B) position of wilting leaves during the hottest parts of the day. Different letters indicate significant differences at P < 0.05 (LSD test). Normal letters indicate the LSD of the interaction (Treatment x Irrigation). Values represent the means ± standard deviations (SD) obtained from 4 plants.

**Supplemental Figure 6:**
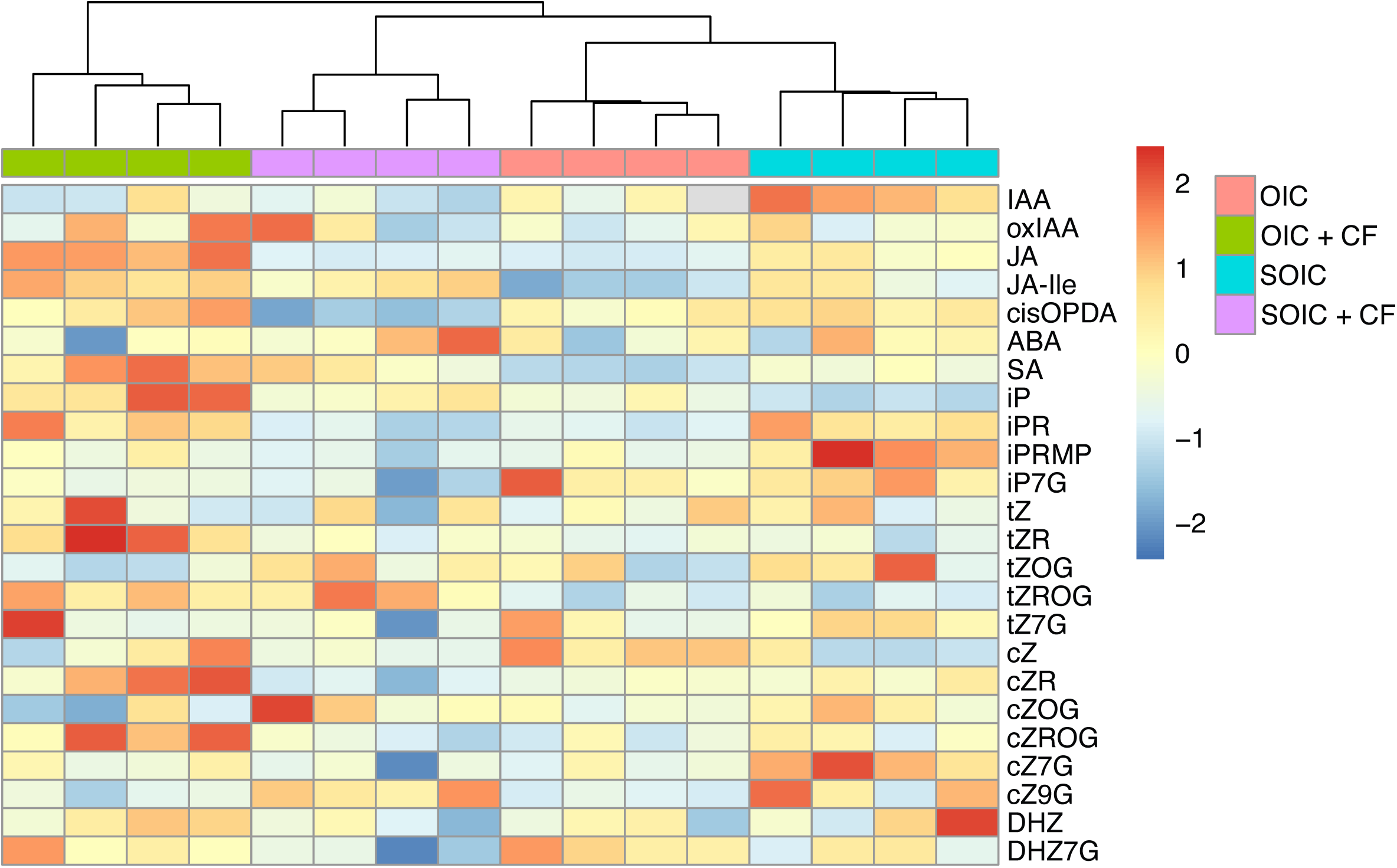
Clustered heatmap of the correlation between treatment and hormone content in leaves of tomato plants grown under optimal and suboptimal irrigation conditions (OIC and SOIC, respectively) with or without the CF treatment.

**Supplemental Figure 7:**
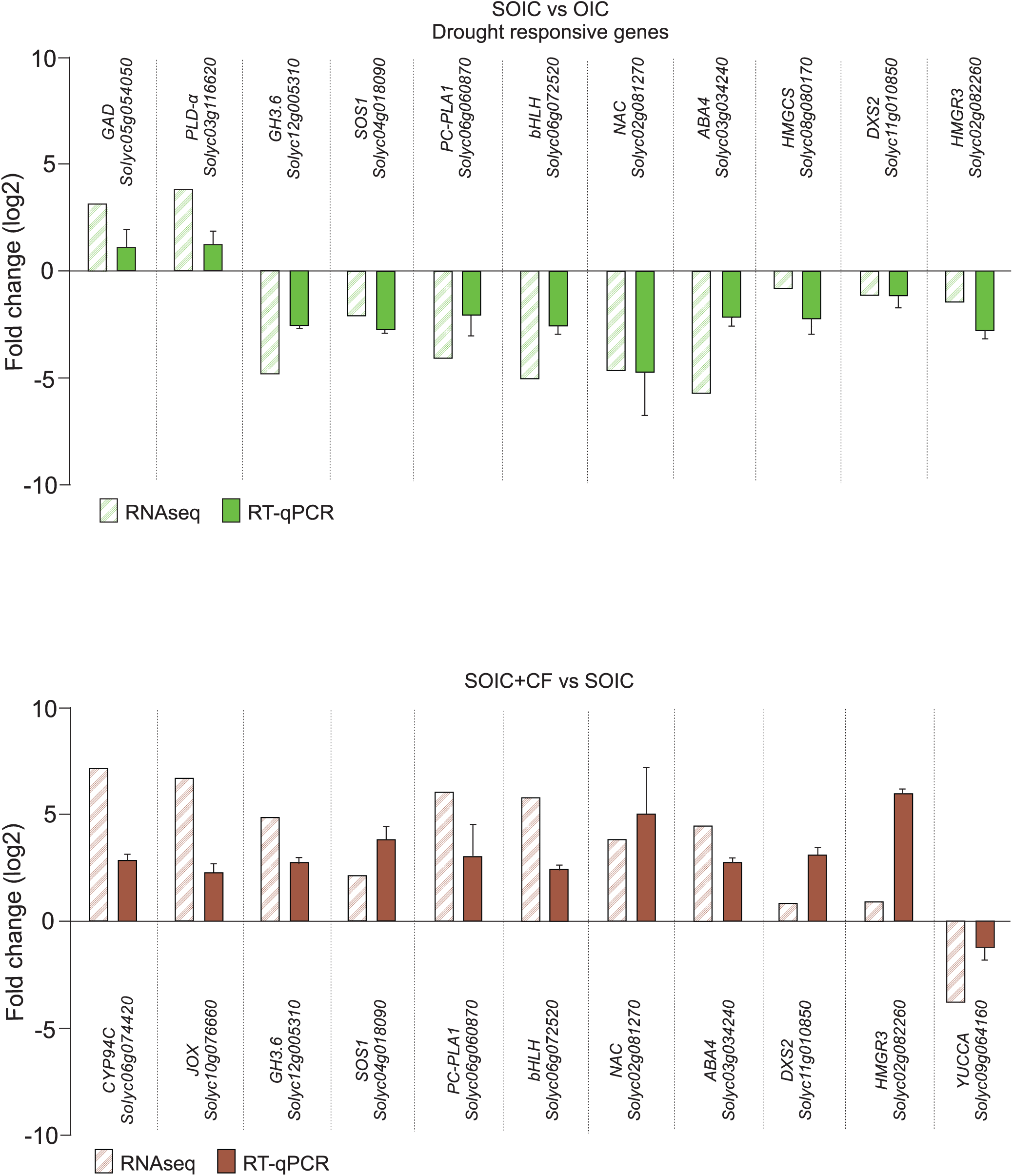
RT-qPCR analysis of selected genes differentially expressed in leaves of tomato plants grown under optimal and suboptimal irrigation conditions (OIC and SOIC, respectively) with or without foliar application of fungal D3 CF dilution.

**Supplemental Figure 8:**
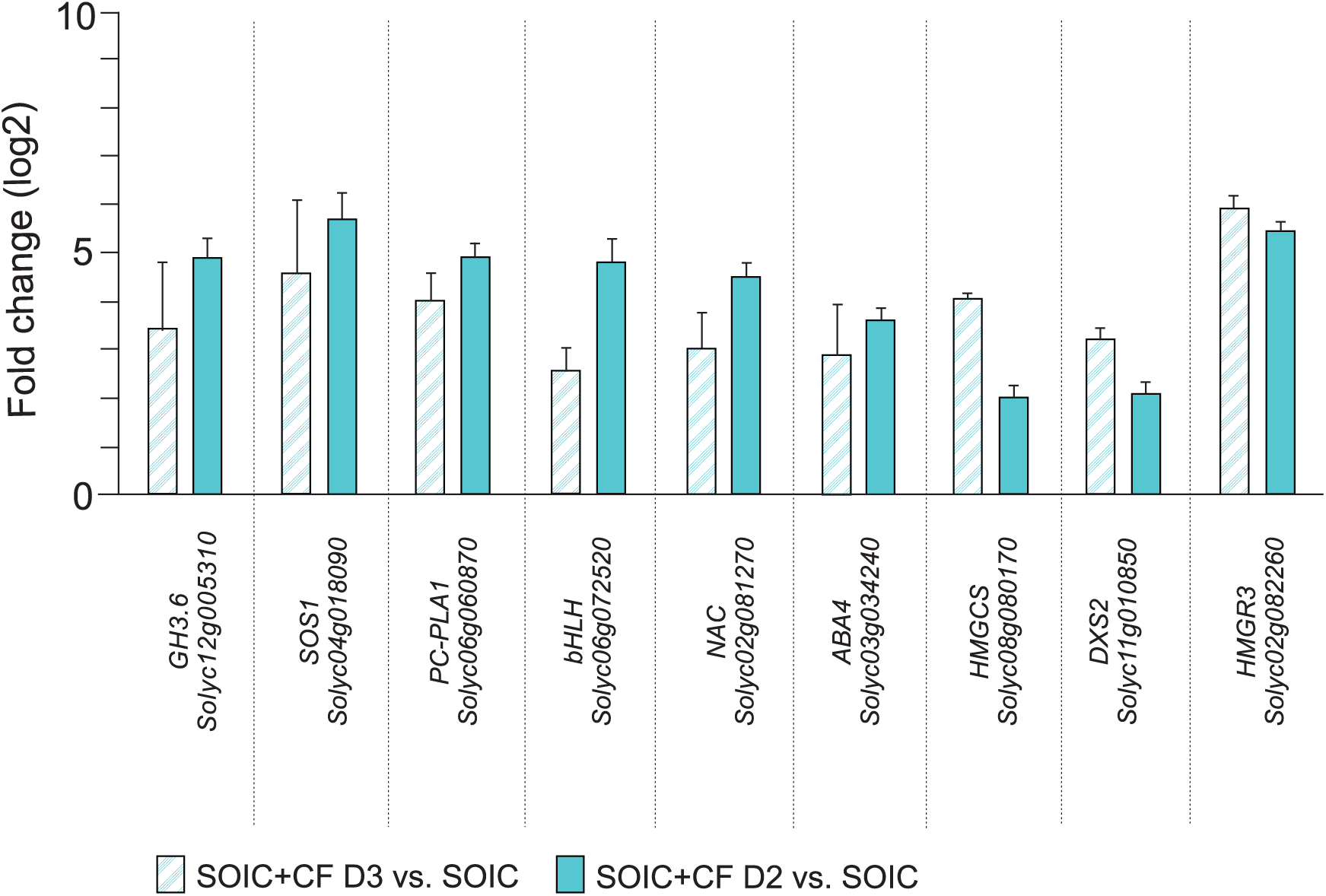
RT-qPCR analysis of selected genes differentially expressed in leaves of tomato plants grown under SOIC with or without foliar application of fungal D2 or D3 CF dilutions.

**Supplemental Figure 9:**
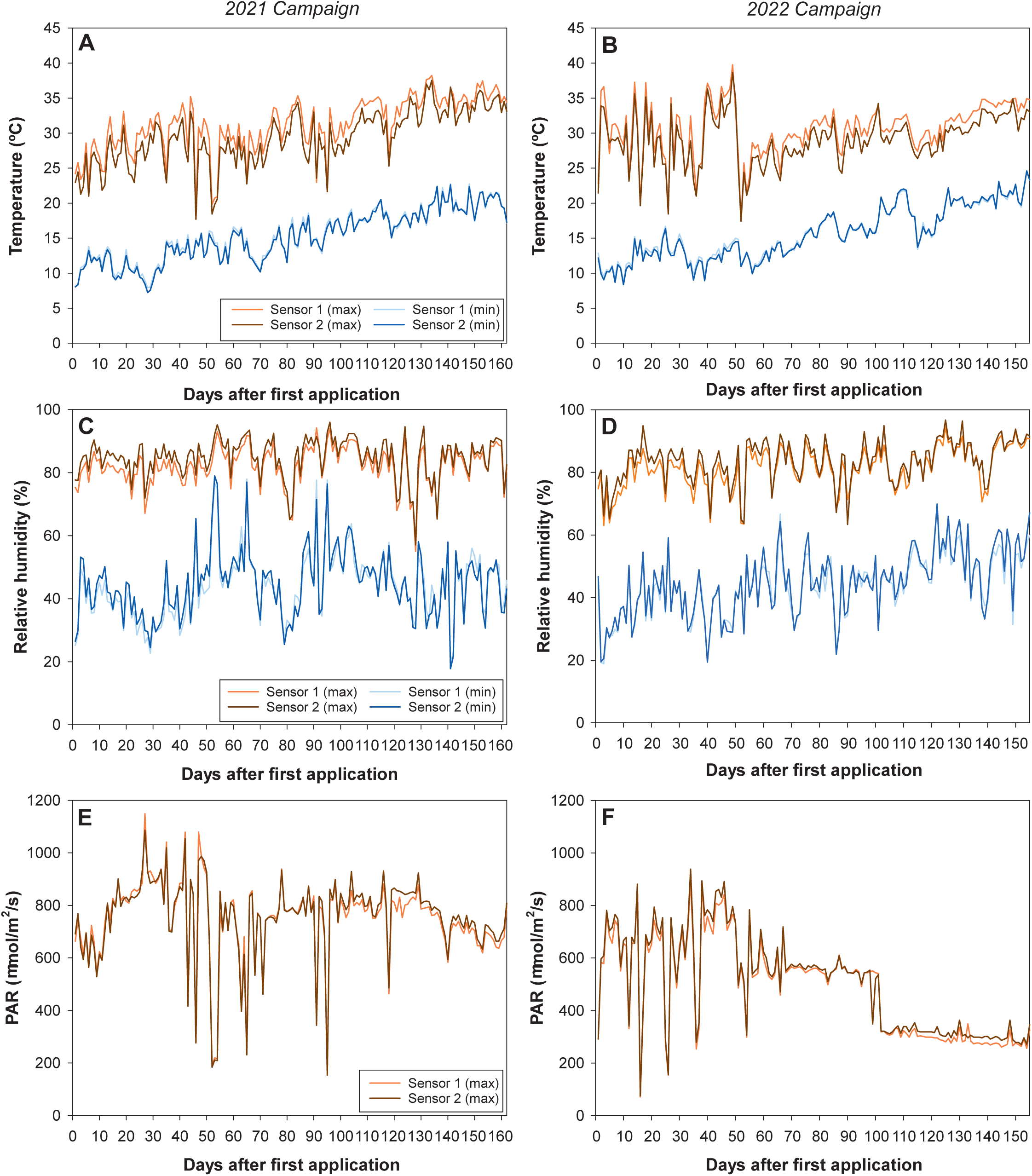
Climatic conditions during the 2021 (A, C, E) and 2022 (B, D, F) campaigns. Maximum (max) and minimum (min) temperatures (A-B), relative humidity (C-D) and maximum photosynthetically active radiation (PAR; E-F) were recorded daily by two sensors.

**Supplemental Figure 10:**
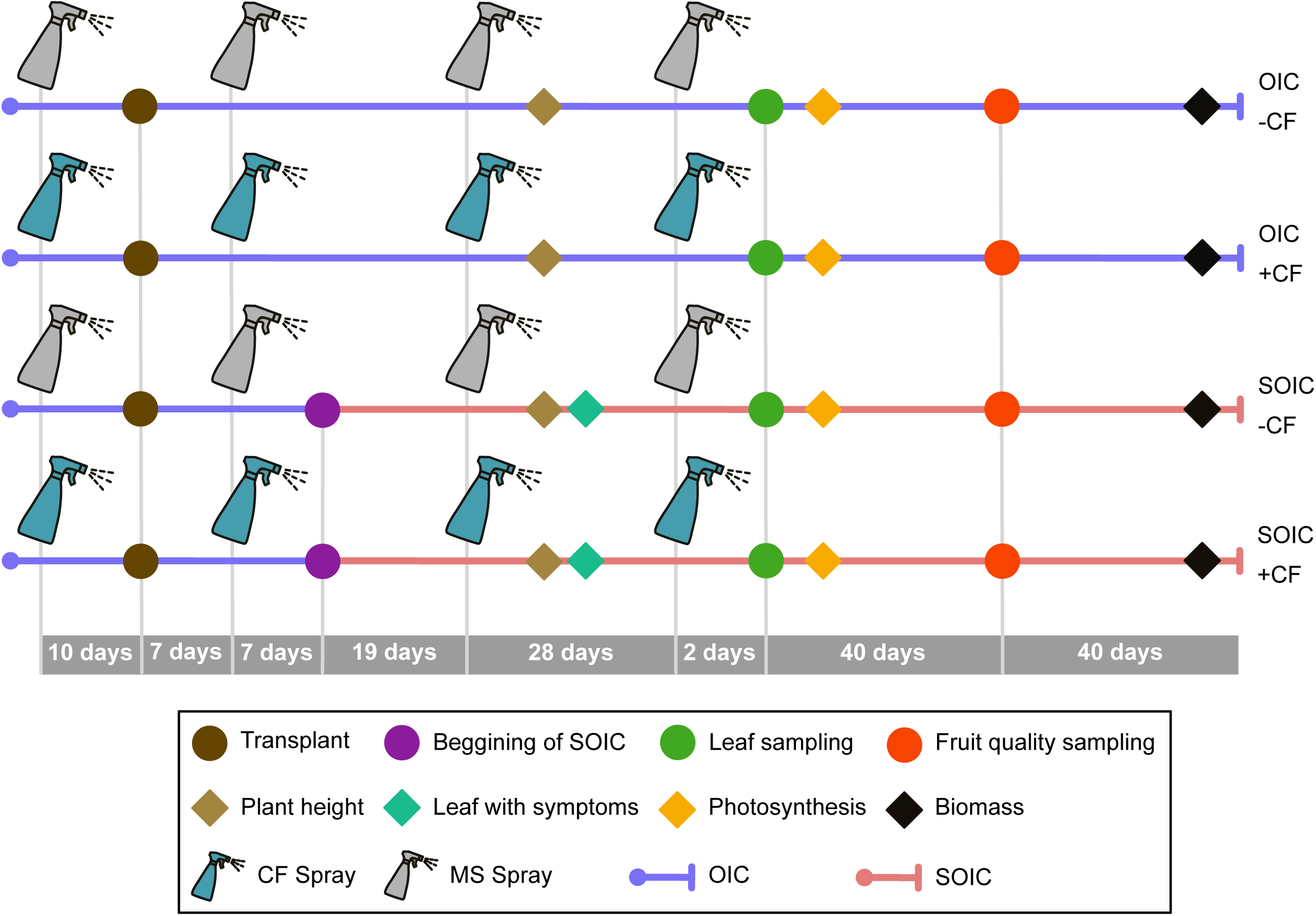
Schematic representation of the experimental design. At the beginning, all plants were irrigated under optimal irrigation conditions (OIC). When seedlings had developed 2-4 true leaves, they were transplanted in 17 L pots filled with a mixture of peat: coconut fibre: vermiculite (45:45:10), and grown under OIC. Fourteen days after transplanting, at the stage of 6-8 true leaves, tomato plants were divided in two groups: plants grown under OIC and plants grown under suboptimal irrigation conditions (SOIC, 60% of OIC irrigation). Foliar applications of diluted (1:6) Murashige and Skoog (MS) medium supplemented with 1.5 mM each of glucose and fructose and CF were made at four moments: 10 days before transplanting, and 7, 33 and 61 days after transplanting. Leaf and fruit materials for biochemical and molecular characterizations were collected 2 and 42 days after the last CF treatment (49 and 89 days after water shortage, respectively).

**Supplemental Figure 11:**
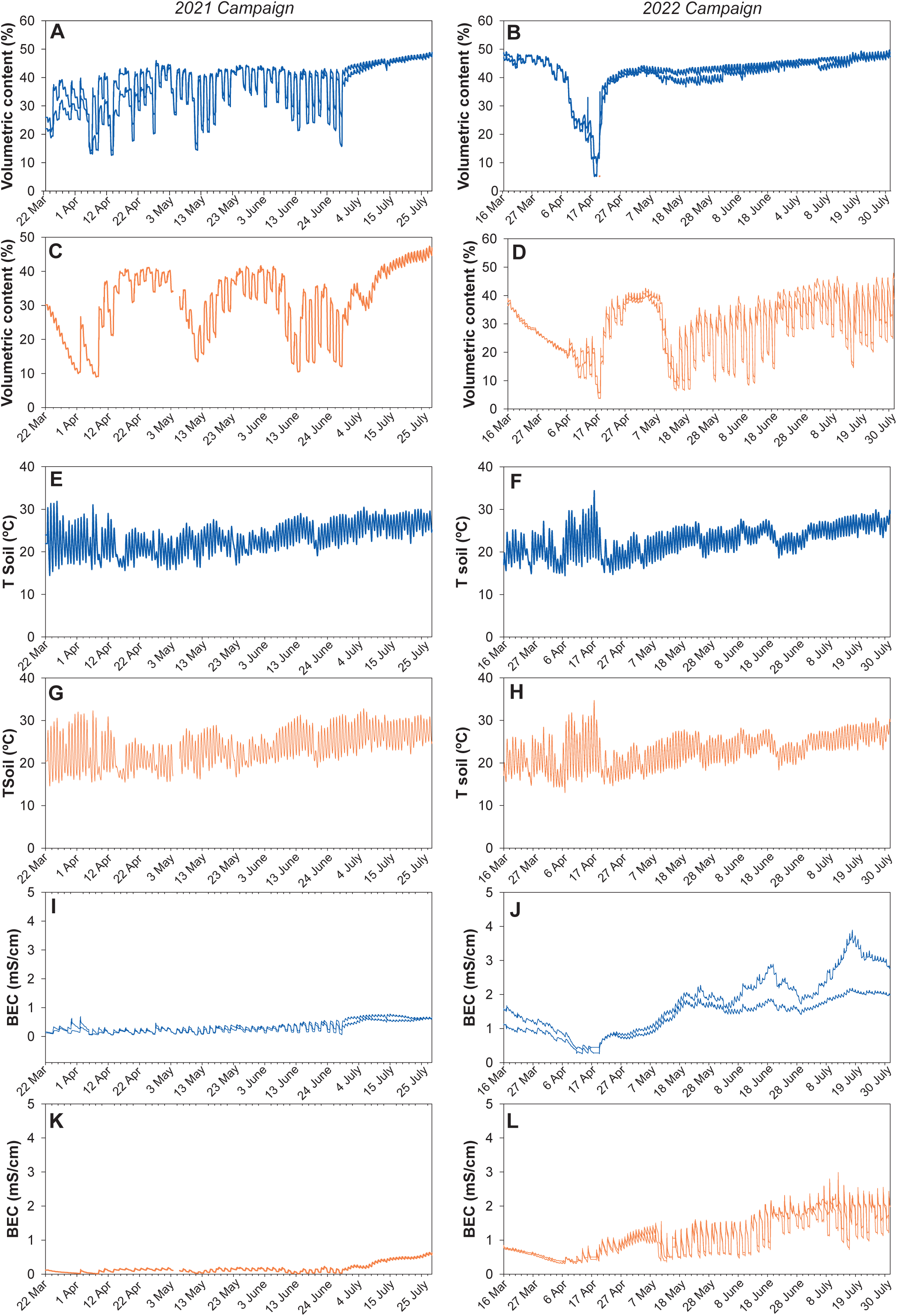
Volumetric content. (A-D), soil temperature (T soil; E-H) and bulk electrical conductivity (BEC; I-L) under optimal irrigation conditions (OIC; A,B) and suboptimal irrigation conditions (SOIC; C-D) during the 2021 (A, C, E, G, I, K) and 2022 (B, D, F, H, J, L) campaigns. In the case of OIC, two sensors were place in two different plants. In the case of SOIC, one sensor was placed.

